# M1 Macrophage-Derived Small Extracellular Vesicles as Synergistic Nanotherapeutics: Harnessing Intrinsic Anticancer Activity and Drug Delivery Capacity

**DOI:** 10.1101/2025.10.21.683755

**Authors:** Gaeun Kim, Hyunsu Jeon, Adrian Chao, James Johnston, Runyao Zhu, Courtney Khong, Yichen Liu, Minzhi Liang, Xin Lu, Yichun Wang

**Author notes:** **Corresponding Author** Yichun Wang - Department of Chemical and Biomolecular Engineering, University of Notre Dame, Notre Dame, Indiana 46556, United States; Harper Cancer Research Institute, University of Notre Dame, Notre Dame, IN 46556, United States.

## Abstract

Small extracellular vesicles (sEVs) have emerged as next-generation multifunctional nanotherapeutics due to their parental-cell traits and role in intercellular communication. Among them, immune cell–derived sEVs are uniquely positioned to couple innate immunomodulatory activities with therapeutic payload delivery, making them highly attractive for cancer therapy. In particular, M1 macrophage-derived sEVs (M1-sEVs) preserve the tumor-suppressive functions of their parent cells, including tumor microenvironment (TME) reprogramming, immune activation, and inhibition of cancer progression. However, the mechanisms by which these activities are coordinated within the TME, and whether they act independently or synergistically, remain poorly understood. Clarifying these mechanisms is crucial for harnessing their intrinsic bioactivity in combination with their natural capacity as drug delivery nanocarriers to optimize therapeutic efficacy. Here, we demonstrate that M1-sEVs exhibit intrinsic stability and circulation longevity via ‘do not eat me’ ligands, as well as tumor-homing ability revealed by proteomic profiling, enabling efficient uptake and deep infiltration in breast cancer models. Functionally, M1-sEVs deliver antiproliferative microRNAs that suppress tumor metabolism, growth, and progression by inhibiting self-renewal, adhesion, migration, motility, and invasion. Importantly, by integrating this endogenous bioactivity with exogenous doxorubicin loading, we achieved synergistic efficacy: a 3-fold reduction in IC_50_ *in vitro* (0.46 μM vs. 1.45 μM for free drug) and 70.18% tumor growth inhibition *in vivo*. These findings highlight M1-sEVs as dual-action nanotherapeutics that combine innate immune-regulatory and tumor-inhibitory functions with efficient drug delivery, advancing their development as powerful platforms for cancer therapy.

## 1. Introduction

Small extracellular vesicles (sEVs) are lipid nanoparticles, in diameter of 50 to 150 nm, naturally secreted by most eukaryotic cells.(1) These nano-sized vesicles carry wide range of cargo, including proteins, lipids, and nucleic acids, inherited from their cells of origin.(1) Their ability to encapsulate and protect these biomolecules while preserving structural integrity and physiological stability during systemic transport further enhances their appeal as nanocarriers for drug delivery.(2) Hence, sEVs have shown promising outcomes in various preclinical studies, and several clinical trials are currently underway.(3–5) Recent studies have shown that sEVs inherit characteristic cargos and membrane proteins, allowing them to preserve the distinctive properties and functions of their parent cells.(6,7) As a result, they can engage in specific biointerface interactions within biological systems, such as targeted recognition and enhanced uptake by their parent cells.(7,8) Specifically, ongoing research aims to harness these inherent, cell-of-origin–dependent attributes in conjunction with advanced engineering strategies to augment sEV functionality, thereby facilitating the development of next-generation multifunctional sEV-based therapeutics.(5,8,9)

sEVs derived from immune cells (e.g., dendritic cells, natural killer cells, and macrophages) have recently been found to play pivotal roles in regulating immune processes such as host defense, inflammation, autoimmune responses, tissue remodeling, and cancer immunity.(10,11) Among all, sEVs derived from M1-polarized macrophages (M1-sEVs) have emerged as promising modulators in cancer research, owing to their ability to carry pro-inflammatory cytokines and chemokines that reprogram the tumor microenvironment, enhance immune responses, and suppress tumor progression.(12) This therapeutic potential stems from the inherent properties of their parent M1 macrophages, the central innate immune cells involved in tumor suppression.(13) For instances, Choo et al. demonstrated that nanovesicles derived from M1 macrophages can improve the efficacy of immune checkpoint inhibitors by reprogramming anti-inflammatory tumor-associated macrophages.(14) Additionally, Wang et al.(15) and Yan et al.(16) reported that M1-sEVs suppress the progression of hepatocellular carcinoma and glioma, respectively. Another study, reported by Wang et al., reported that M1-sEVs promote apoptosis in lung adenocarcinoma cells.(17) While M1-sEVs stand out for their use in cancer therapeutics, with the capacity to modulate immune responses, regulate inflammatory pathways, and exert direct anti-tumor activity, the mechanisms by which these functions are coordinated within the tumor and its microenvironment (TME) and whether they can be harnessed to optimize therapeutic efficacy have not yet been fully explored.(12,13) Thus, it is essential to gain a deeper understanding of how M1-sEVs operate through distinct, parallel effects or through coordinated and synergistic mechanisms, since this distinction carries significant implications for their therapeutic application in cancer. Moreover, clarifying how these mechanisms can be harnessed in conjunction with their promising capacity as nanocarriers will be critical for optimizing drug delivery strategies.

Herein, we investigate how the intrinsic antitumor functions of M1-sEVs can be strategically harnessed to maximize their synergistic therapeutic potential while simultaneously enabling efficient chemotherapeutic drug delivery, thereby establishing them as dual-role nanomedicines for effective cancer treatment (Figure 1). We first demonstrated that M1-sEVs exhibit low immunogenicity by expressing the CD47 ligand on their surface, enabling them to evade immune clearance and prolong systemic circulation.(18) This potentially complements their additional tumor-homing function, as we revealed the expression of targeting ligands and tumor microenvironment (TME)-recruiting proteins (β2 integrin, Gal-3, VEGFR1, CD44, and NRP1) on M1-sEVs, which, in turn, facilitated enhanced cellular uptake and deeper penetration in solid tumors. Additionally, we identified specific miRNAs (miR-150, miR-181a, and miR-34a) within M1-sEVs that are associated with the suppression of cancer cell proliferation, which was further supported by significant inhibition of cell growth, impaired scratch closure, and reduced colony formation. Building on these inherent biological advantages, we further utilized M1-sEVs as drug delivery vehicles by loading them with the chemotherapeutic agent doxorubicin (Dox). By employing our previously developed chiral graphene quantum dot (GQD)-assisted loading strategy,(19) we achieved efficient Dox encapsulation within M1-sEVs, with a loading efficiency of 63.37 ± 2.65%. *In vitro*, Dox-loaded M1-sEVs reduced the IC_50_ in MCF-7 cells from 1.45 μM (free Dox) to 0.46 μM after 48 hours, demonstrating significantly increased cytotoxicity through synergistic delivery. *In vivo* studies further validated this effect, with Dox-loaded M1-sEVs (Dox/M1-sEV) achieving substantial tumor growth inhibition (70.18 ± 11.37%). This overall strategy achieved a potent synergistic therapeutic effect by combining the intrinsic anticancer activity of M1-sEVs with chemotherapeutic delivery, resulting in markedly enhanced efficacy against breast cancer. Collectively, these findings highlight a robust therapeutic platform that leverages both the inherent antitumor functions of M1-sEVs and chemotherapy delivery capacity to substantially improve treatment outcomes, underscoring its potential impact in advancing next-generation cancer therapeutics.

**Figure 1.**
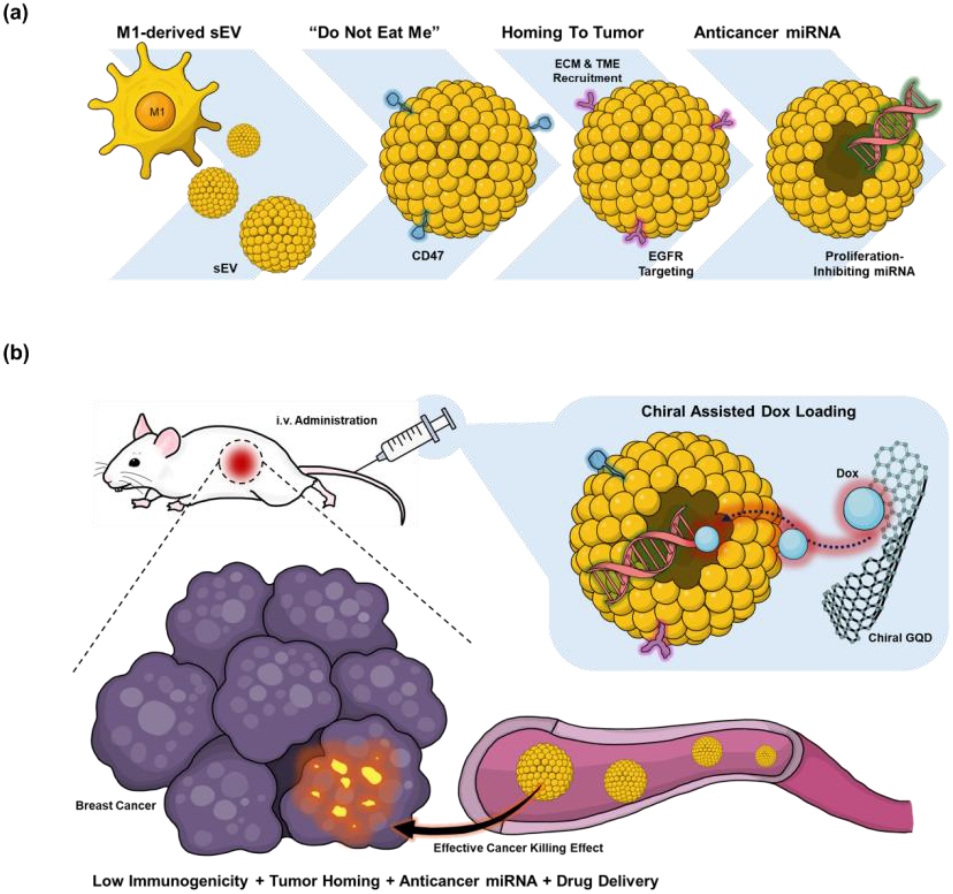
Schematic illustration of (a) the intrinsic tumor-suppressive potential of M1-derived small extracellular vesicles (M1-sEVs), and (b) the synergistic therapeutic strategy combining M1-sEVs with chemotherapeutic payloads for enhanced anti-cancer efficacy.

## 2. MATERIALS AND METHODS

### 2.1 Materials

Carbon nanofibers (719803), Hydrochloric acid (320331), BioTracker 488 Green Nuclear Dye (SCT120), PKH26 (MINI26-1KT), Alginic acid sodium salt (180947), calcium chloride dihydrate (CaCl2; 223506), sodium hydroxide anhydrous (NaOH; S5881), and alginate lyase powder (A1603) were purchased from Sigma-Aldrich (MO, USA). Sulfuric acid (BDH3068-500MLP), Nitric acid (BDH3044-500MLPC), syringe filter (0.22 μm; 76479-010), Fetal bovine serum (FBS; 1300-500), HyClone Dulbeccos Modified Eagles Medium with high glucose (DMEM/HIGH GLUCOSE; 16750-074), SylgardTM 184 silicone elastomer (polydimethylsiloxane, PDMS; 102092-312) were purchased from VWR (PA, USA). Dialysis membrane tubing (MWCO: 1 kD; 20060186) was purchased from Spectrum Chemical Manufacturing Company (NJ, USA). 1-ethyl-3-(3-dimethyl-aminopropyl) carbodiimide (EDC; 22980), VybrantTM Multicolor Cell-Labeling Kit (DiO, DiD; V22889), Pierce BCA Protein Assay Kits (23225), GibcoTM Trypsin-EDTA (25200072), Prestoblue Cell Viability Reagent (A13261), eBioscience Lipopolysaccharide Solution (LPS; 500X; 00-4976-93), anti-CD86 (B7-2) antibody (12-0862-82), anti-CD206 (MMR) antibody (12-2061-82), Mouse IL-6 Uncoated ELISA Kit (88-7064-88), and LIVE/DEAD™ Cell Imaging Kit (488/570) (R37601) were purchased from Thermofisher Scientific (MA, USA). N-hydroxysuccinimide sodium salt (Sulfo-NHS; 56485), and D-cysteine (A110205-011) were purchased from AmBeed (IL, USA). Sodium Hydroxide (1310-73-2) was purchased from Ward’s Science (NY, USA). Nitrocellulose membrane (1662807), and Clarity Max Western Enhanced Chemiluminescence (ECL) Substrate (1705060) were purchased from Bio-Rad (CA, USA). RIPA Buffer (9806), anti-mouse Horseradish Peroxidase (HRP)-linked secondary antibody (7076) were purchased from Cell Signaling Technology (MA, USA). Anti-beta-actin antibody (sc-47778), Anti-CD9 antibody (sc-13118), anti-CD63 antibody (sc-5275), anti-CD81 antibody (sc-166029), anti-CD47 antibody (sc-53050) were purchased from Santa Cruz Biotechnology (TX, USA). Phosphate-buffered saline (PBS; 21-040-CM) was purchased from Corning (NY, USA). Eagle’s Minimum Essential Medium with L-glutamine (EMEM; 30-2003) was purchased from American Type Culture Collection (ATCC; VA, USA). Antibiotic antimycotic (15240096) was purchased from Fisher Scientific (MA, USA). 4% Paraformaldehyde (PFA; 15735-50S), Aqueous Glutaraldehyde 25% Solution Biological Grade (16400), Crystal Violet 1% Aqueous (26088-10), and UranyLess (22409) were purchased from Electron Microscopy Sciences (PA, USA). Tween-20 (BTNM-0080) was purchased from G-Biosciences (MO, USA). Cell Counting Kit-8 (CCK-8; ALX-850-039-KI01), and Doxorubicin.HCl (BML-GR319-0025) were purchased from Enzo Biochem (NY, USA). Cryopres dimethyl sulfoxide (DMSO; 092780148) was purchased from MP Biomedicals (CA, USA). 5-[(4,6-Dichlorotriazin-2-yl)amino]fluorescein hydrochloride (5-DTAF; 21811-74-5) was purchased from Chemodex (Switzerland). Cell ExplorerTM Live Cell Tracking Kit (22620) was purchased from AAT Bioquest (CA, USA). miRNA All-In-One cDNA Synthesis Kit (G898), BlasTaq™ 2X qPCR MasterMix (G891), U6 Primers (MPH00001), mmu-miR-150-5p Primers (MPM00868), mmu-miR-150-3p Primers (MPM00867), mmu-miR-181a-5p Primers (MPM00889), mmu-miR-34a-5p Primers (MPM01238), and mmu-miR-34a-3p Primers (MPM01237) were purchased from Applied Biological Materials Inc (BC, Canada).

### 2.2 Instruments

Transmission electron microscopy (TEM; Talos F200i (S); Thermofisher Scientific, MA, USA) was used to confirm the nanostructures. Nanoparticle tracking analysis (NTA; NanoSight LM10 system; Malvern Instrument Ltd., Worcestershire, UK) was used to measure the size distribution and concentration of nanoparticles. Two-photon confocal microscopic (TP-CLSM; Leica Stellaris 8 DIVE Point Scanning Confocal Microscope; Leica Microsystems, Wetzlar, Germany) was used to image 3D spheroid models. Single-molecule localization microscopy (SMLM; ONI, CA, USA) was used for super-resolution microscopic images of individual sEVs. Fluorescence was measured by a plate reader (Infinite 200 PRO; Tecan, Männedorf, Switzerland). Circular dichroism (CD) spectrometer (Jasco J-1700 Spectrometer; Jasco International Company, MD, USA) was used to measure the absorbance of polarized light were measured. Bruker Tensor 27 Fourier-transform infrared (FTIR) Spectrometer (Bruker Optics International Company, MA, USA) was used to analyze the Attenuated Total Reflectance (ATR)-FTIR. Mass spectrometer (MS; Thermo Q-Exactive HF, Thermo Fisher, MA, USA) was used to identify the proteins. Western blot and agarose gel images acquired using two bioimaging systems (Azure 600; Azure Biosystems, CA, USA). Humidified incubator (MCO-15AC; Osaka, Japan) was used to maintain and incubate all cell lines used in this study. Fluorescence-activated cell sorting system (FACS; Cytek Northern LightsTM (NL)-CLC NL-2000; Cytek Biosciences, CA, USA) was used to sort and detect the cells. FACS data was analyzed using FlowJo software v.10.10.0 (FlowJo LLC, OR, USA). Advanced Molecular Imaging HT Instrument (SPECTRAL AMI HT; Spectral Instruments Imaging, AZ, USA) was used for *ex vivo* mice study. CFX Connect Real-Time PCR Detection System (Bio-Rad, CA, USA) was used for quantitative reverse transcription polymerase chain reaction (qRT-PCR). Brightfield microscope (Keyence BZ-X810; Keyence Corporation, IL, USA) was used for H&E histological imaging. SigmaPlot 10.0 (Systat Software Inc., CA, USA) was used to generate all plots of the analyzed data.

### 2.3 Macrophage Differentiation and sEV Isolation

Undifferentiated RAW 264.7 murine macrophage cells were cultured in high-glucose DMEM supplemented with 10% FBS and 1% antibiotic-antimycotic in a humidified incubator at 37°C with 5% CO2. The cells were washed with 1× PBS, trypsinized with Trypsin-EDTA for passages, and incubated for at least 1 day before any experiment. RAW 264.7 cells were stimulated with LPS to induce differentiation into M1-type macrophages.(20) Successful M1 polarization was confirmed by FACS using anti-CD86 and anti-CD206 antibodies conjugated with a fluorophore. Once mouse fibroblast cells (3T3), RAW 264.7 cells (M0), and LPS-stimulated RAW 264.7 cells (M1) reached 70–80% confluency in the flasks, the culture media were replaced with serum-free media, followed by three washes with 1× PBS. After 24 hours of incubation, the serum-free media from all three cell types were collected and processed using vacuum filtration systems (0.22 μm pore size; 10040-460; VWR, PA, USA) to remove undesired large debris. The filtered media was then subjected to ultrafiltration using centrifugal devices (100 kDa cutoff; Spin-X UF Concentrator; Corning, NY, USA) and washed with 1× PBS buffer until transparent 2 mL sEV suspension was collected. The nanostructure of sEVs was observed under TEM with negative staining by UranlyLess. NTA measurements were performed using diluted sEVs (100 times dilution; 10 μL in 1 mL of PBS buffer) and analyzed with the NTA 3.3 analytical software suite. The expression of sEV markers (CD9, CD63, and CD81) and the immune evasion marker CD47 were confirmed by Western blot analysis.

### 2.4 In Vivo Safety Evaluation of sEVs

Total of 1.5 × 10^9^ sEVs suspended in 200 µL PBS were intravenously administered to mice (female, C57BL/6) in four groups: Ctrl, 3T3-sEV, M0-sEV, and M1-sEV. The Ctrl group was administered 200 µL of PBS without sEVs per mouse. Body weight of the mice was monitored every 2 days for 2 weeks following injection (Figure S4a). 2 weeks post-injection, blood samples were collected, and serum was isolated for IL-6 quantification using an enzyme-linked immunosorbent assay (ELISA) (Figure S4b). At the same time point (2 weeks post-injection), major organs (brain, heart, liver, spleen, kidney, and lung) and muscle were harvested, paraffin-embedded, and sectioned for histological analysis using hematoxylin and eosin (H&E) staining (Figure S4c).

### 2.5 *Ex Vivo* Biodistribution Study of sEVs

sEVs were labeled with the lipophilic dye DiD and washed three times with PBS to remove any unincorporated dye. A total of 1.5 × 10^9^ sEVs suspended in 200 µL PBS were intravenously administered to mice (female, C57BL/6) divided into 4 groups: Ctrl, 3T3-sEV, M0-sEV, and M1-sEV. The Ctrl group received 200 µL of PBS without sEVs. Mice were sacrificed at 1, 3, and 6 hours post-injection, and major organs (brain, heart, liver, spleen, kidney) as well as muscle were harvested for *ex vivo* fluorescence imaging. Imaging analysis was performed using Aura Imaging Software v4.5 (Spectral Instruments Imaging, AZ, USA) and ImageJ software. All samples were initially measured in a black 96-well plate prior to administration (Figure S3a), and all *ex vivo* fluorescence data were normalized to the signal of the 3T3-sEV samples for comparative quantification.

### 2.6 Proteomic and Transcriptomics Characterization of sEVs

Western blot analysis was performed as follows: sEVs were lysed using 1× RIPA buffer, and total protein concentrations were determined with the BCA assay. Equal amounts of protein (15 μg per sample) were separated by SDS-PAGE and transferred onto nitrocellulose membranes. The membranes were probed with primary antibodies against CD9, CD63, CD81, CD47, and β-actin. After incubation with HRP-conjugated secondary antibodies, immunoreactive bands were visualized using an ECL substrate and imaged with a bioimaging system. Sample preparation for MS-based proteomics was conducted as follows: cell-derived sEVs were lysed using 1× RIPA buffer, and 50 μg of total protein was subjected to trypsin digestion. The resulting peptides were eluted, dried, and resuspended before being loaded onto an ultrahigh-resolution mass spectrometer coupled with a nano-ultrahigh-pressure liquid chromatography (nano-UHPLC) system for bottom-up proteomic analysis. GraphPad Prism 8.0 (GraphPad Software Inc., CA, USA) was used to generate tables for barcode plots to visualize protein identification. MicroRNA (miRNA) identification was performed as follows: RNA was extracted from sEVs using the Exosomal RNA Isolation Kit (58000; Norgen Biotek Corp, ON, Canada). The extracted RNA was then confirmed the quality control (QC) using the Qubit assay. Followed by, qRT-PCR was performed as follows: RNA isolated from sEVs was reverse transcribed using the miRNA All-In-One cDNA Synthesis Kit. qRT-PCR was performed using BlasTaq™ 2X qPCR MasterMix on CFX Connect Real-Time PCR Detection System. Amplification was carried out at 95°C for 3 minutes, followed by 40 cycles of 15 seconds at 95°C, and 1 minute at 60°C. Gene-specific primers were used, with U6 small nuclear RNA (snRNA) serving as an internal housekeeping control for miRNA expression analysis. The data is presented as relative changes in gene expression with respect to the 3T3-sEV sample.

### 2.7 Cancer Cell Uptake and Infiltration of sEVs

FACS analysis was performed as follows: MCF-7 cells were seeded at 5 × 10^5^ cells per well in a 24-well plate. sEVs were labeled with the lipophilic dye DiD and washed three times with PBS to remove unincorporated dye. DiD-labeled sEVs were then co-incubated with MCF-7 cells at a concentration of 1 × 10^3^ sEVs per cell. For the time-dependent uptake study, cells were incubated with DiD-sEVs for 1, 4, and 8 hours at 37 °C. For the temperature-dependent uptake study, cells were incubated with DiD-sEVs for 4 hours at 4 °C. After incubation, DiD-sEV positive cells were quantified by FACS. Cell spheroid generation in a hydrogel framework was performed as follows: An inverted colloidal crystal (iCC) framework using low-melting agarose from a previous study(21) was employed to form MCF-7 spheroids. To label LM agarose with 5-DTAF, agarose was hydrated and reacted with a 10 mM 5-DTAF solution in DMSO and 1 M NaOH in DI water under basic conditions, followed by multiple washing and centrifugation steps to isolate the labeled agarose. The labeled agarose was then blended with unlabeled hydrated agarose at a 1:9 weight ratio, cast, thermally gelled, and reshaped into desired forms using controlled heating and cooling cycles. Alginate microgels (e.g., 2.5%w/v, 250 µm) were packed via sonication and templated in a 5-DTAF-tagged agarose hydrogel (e.g., 15.6%w/v), with alginate subsequently degraded using alginate lyase and PBS. MCF-7 cells, at >80% confluency, were trypsinized, stained with 20 µM Cell ExplorerTM blue live cell tracker, and condensed to 1 × 10^7^ cells/mL for seeding. The previous centrifugal method(21) was used to seed 5 × 10^6^ cells per run, totaling 1 × 10^7^ cells per framework. The culture media was refreshed in 72 hours after cell seeding into the framework while further refreshed in every 48 hours. The sEV delivery evaluation for MCF-7 spheroids was performed as follows: For evaluating sEV delivery/penetration in MCF-7 spheroids in the iCC framework, total four sample groups were designated: (1) Control (Ctrl; PBS), (2) 3T3-sEV, (3) M0-sEV, and (4) M1-sEV. All sEVs were labeled with the lipophilic dye DiD and washed three times with PBS to remove any unincorporated dye. After 144 hours post-seeding and PBS washing, MCF-7 spheroid frameworks were treated with each sEVs, at an amount of 4.8 × 10^8^ in 2 mL of MCF-7 media. After 24 hours from sEVs treatment, all samples were washed with PBS and immersed in 2 mL of PBS after being fixed with 2% PFA for 3 h. The live cell signal (i.e., fluorescence intensity; FILive Cell) and sEV signal (i.e., FIsEV) from spheroid regions were collected under multi-photon imaging from a point scanning confocal microscope (TP-CLSM; Leica Stellaris 8 DIVE, Leica; Germany). To quantitatively analyze fluorescent signals from sEV-treated spheroids within the framework, we implemented an automated Python-based image processing method for TP-CLSM images, building on previous work(22) which is available on Github (https://github.com/HyunsuJeon-ND/iCC_Automated_HCHT_Radial_Analysis). Multiphoton z-stack images from iCC frameworks were split into grayscale fluorescence channels, with circular ROIs identified via OpenCV-based Hough Circle Detection and uniformly applied across channels. Spheroid boundaries were defined using composite masks derived from thresholded red and blue channels, and centroids were calculated by identifying the maximal cross-sectional Z-plane and computing fluorescence-weighted XY positions; radial signal intensities were then extracted for each spheroid to generate spatial distribution curves, penetration profiles.

### 2.8 Anti-Proliferative Effects of M1-sEVs on Cancer Cells

WST-8 and Resazurin assays were performed as follows (Figure S15a): MCF-7 cells were seeded in transparent 96-well plates for the WST-8 assay and black 96-well plates for the Resazurin assay at a density of 1 × 10^4^ cells per well. After 24 hours, sEVs were added at a concentration of 1 × 10^3^ sEVs per cell and co-incubated at 37 °C. At 1, 2, 3, and 6 days post-treatment, WST-8 reagent (from the CCK-8 kit) and Resazurin reagent (from the PrestoBlue kit) were added. Absorbance (460 nm) for WST-8 and fluorescence (Ex/Em: 510/610 nm) for Resazurin were measured using a microplate reader. Scratch assay was conducted as follows (Figure S15b): MCF-7 cells were seeded in transparent 6-well plates at a density of 5 × 10^4^ cells per well. After 72 hours, 5 × 10^3^ sEVs per cell were added and co-incubated for 24 hours at 37°C. Subsequently, a sterile 20 µL pipette tip was used to create a linear scratch in each well. Scratch closure was monitored at 0, 24, and 48 hours post-scratch using a light microscope. The scratched areas were quantified using ImageJ software, and the relative wound area was normalized to the untreated control group. Clonogenic assay was conducted as follows (Figure S15c): MCF-7 cells were seeded in transparent 6-well plates at a density of 5 × 10^4^ cells per well. After 24 hours, sEVs were added at a concentration of 1 × 10^3^ sEVs per cell and co-incubated for 72 hours at 37 °C. Following incubation, cells were washed, trypsinized, and re-seeded into 6-well plates at a density of 3 × 10^3^ cells per well. Fresh media was replaced every two days throughout the incubation period. On days 3, 6, and 10, media was aspirated, and cells were washed three times with ice-cold 1× PBS. Colonies were fixed with 6% glutaraldehyde for 5 minutes at room temperature and stained with 0.5% (w/v) crystal violet for 15 minutes. Each well was photographed, and colony numbers were quantified using ImageJ software. Relative colony formation (%) was normalized to the untreated control group.

### 2.9 Chiral GQDs-Assisted Dox Loading into sEVs

The Dox/*D*-GQD complex was characterized by fluorescence quenching analysis:(19) A fixed concentration of doxorubicin (200 μM) was incubated with increasing concentrations of *D*-GQDs (0–22.5 μM). As Dox interacted with *D*-GQDs, fluorescence self-quenching was observed, reaching a plateau at higher *D*-GQD concentrations. Relative quenching was calculated using the following equation:

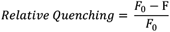

Dox encapsulation efficiency was quantified using a method specifically developed for our chiral GQD loading approach (Figure S14a): First, a calibration curve was first generated to establish the linear relationship between Dox concentration and its fluorescence intensity (excitation/emission: 490/560 nm). Dox fluorescence was measured with excitation at 490 nm and emission at 560 nm. To extract the Dox/*D*-GQD complexes, the lipid membrane of the sEVs was disrupted using Tween-20 (10X dilution), releasing the encapsulated cargo. The extracted Dox/*D*-GQD complexes were then dissolved in acetonitrile (10X dilution) and thoroughly sonicated to disrupt pi stacking interactions mediated by the GQDs, as their intrinsic fluorescence could interfere with accurate measurement of the Dox signal. Subsequently, the mixture was filtered through a 2 kDa Amicon tube to separate the Dox from the *D*-GQDs. The resulting filtrate-eluent containing free Dox was collected, and its fluorescence intensity was used to estimate the encapsulation efficiency using the following equation:

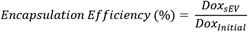

For supportive validation, an additional fluorometric approach, the fluorescence recovery assay,(19) was performed using a plate reader. DiO-stained (ex/em: 470/520 nm) and Dox-loaded (ex/em: 490/560 nm) sEVs were analyzed by comparing fluorescence intensity before and after lysis with Tween-20. Furthermore, SMLM was used to image individual sEVs stained with DiO (green), loaded with Dox (red) and *D*-GQDs (blue) complexes.

### 2.10 Synergistic Anticancer Effects of Dox-Loaded M1-Derived sEV

*In vitro* cell viability was evaluated as follows (Figure S15d): MCF-7 cells were seeded in transparent 96-well plates at a density of 1 × 10^4^ cells/well. After 24 hours, Dox-loaded sEVs were added and incubated at 37 °C. At 24 and 48 hours post-treatment, Cell Counting Kit-8 (CCK-8) reagent was added, and absorbance was measured at 460 nm to determine relative cell viability (%). The results were normalized to the Ctrl group, which received PBS treatment. IC 50 were calculated using the formula:

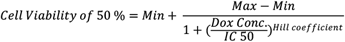

*In vivo* therapeutic efficacy was evaluated as follows (Figure 5e): Orthotopic xenograft mouse models were established by subcutaneously injecting 1 × 10^6^ MCF-7 cells suspended in 100 µL of PBS into the mammary fat pad of mice. (female, NCr nude). 10 days after tumor cell injection, mice were randomly assigned to five treatment groups: Ctrl, Dox, Dox/3T3-sEV, Dox/M0-sEV, and Dox/M1-sEV. The treatment was intravenously administered. The Ctrl group was administered with 150 µL of PBS per mouse, while both the free Dox and Dox-loaded sEV groups were administered 4 mg/kg of Dox (∼150 µL/mouse). Dox concentration in sEVs was calculated based on the loading efficiency. Treatments were administered every two days for a total of three doses. Tumor size was measured on days 0, 2, 4, 7, 10, and 14. The representative tumor images were acquired on each measurement day. Tumor volumes were calculated using the formula:(23)

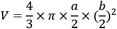

a and b represent the maximum and minimum tumor diameters, respectively. Tumor growth inhibition (TGI) was calculated using the following formula:(23)

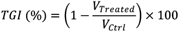

After the treatment period, major organs (brain, heart, liver, spleen, kidney, and lung), muscle, and tumor tissues were collected, paraffin-embedded, sectioned, and stained with H&E for histological imaging analysis.

## 3. RESULTS

### 3.1. Isolation and Comprehensive Characterization of sEVs

Using an optimized LPS dose range of 0–1,000 ng/mL, we successfully generated M1 macrophages from M0 precursors (Figure S1). FACS analysis confirmed expression of the M1 surface marker CD86, with negligible expression of the M2 marker CD206, following 6 h of LPS co-incubation.(20) Based on these results, we selcted 500 ng/mL LPS as the optimal stimulation dose (Figure 2a) for subsequent sEV isolation from M1-sEV. For comparative profiling, we isolated sEV derived from M0 macrophages (M0-sEVs) and 3T3 fibroblasts (3T3-sEVs) using our established size-based ultrafiltration method to ensure consistent yield and purity.(8,24,25) The spherical morphology of sEVs was confirmed by TEM (Figure 2b–d), with size distribution analyses revealing mean diameters of 79.73 ± 24.08 nm for 3T3-sEVs, 82.72 ± 24.61 nm for M0-sEVs, and 91.66 ± 19.16 nm for M1-sEVs. Structural integrity was further supported by NTA (Figure 2f-h), which demonstrated a distinct and narrow size distribution, yielding mean sizes of 133.8 ± 36.2 nm for 3T3-sEVs, 134.6 ± 31.5 nm for M0-sEVs, and 134.4 ± 42.5 nm for M1-sEVs. The smaller sizes of sEVs observed by TEM were potentially attributed to dehydration-induced shrinkage and flattening of vesicles under vacuum, as well as exposure to high-energy electron beams.(26,27) Conversely, NTA tended to analyze slightly larger sizes, as its measurement is based on tracking Brownian motion via light scattering, which limits sensitivity for smaller nanoparticles.(27) Western blot analysis confirmed the presence of standard sEV markers (CD9, CD63, and CD81) on all three types of sEVs. Notably, we identified the immunomodulatory ligand CD47 on both macrophage-derived sEVs (M0-sEVs and M1-sEVs), whereas its expression was minimal on 3T3-sEVs. As a ‘self’ recognition marker, CD47 engages immune cell receptors to deliver a ‘do not eat me’ signal that suppresses macrophage-mediated phagocytosis.(28) Consistently, macrophages themselves express CD47, suggesting a mechanism of self-protection against clearance.(29) Thus, the distinct CD47 expression patterns between macrophage derived sEVs and 3T3-sEVs reflected the inheritance of membrane proteins from their parental cells.

**Figure 2.**
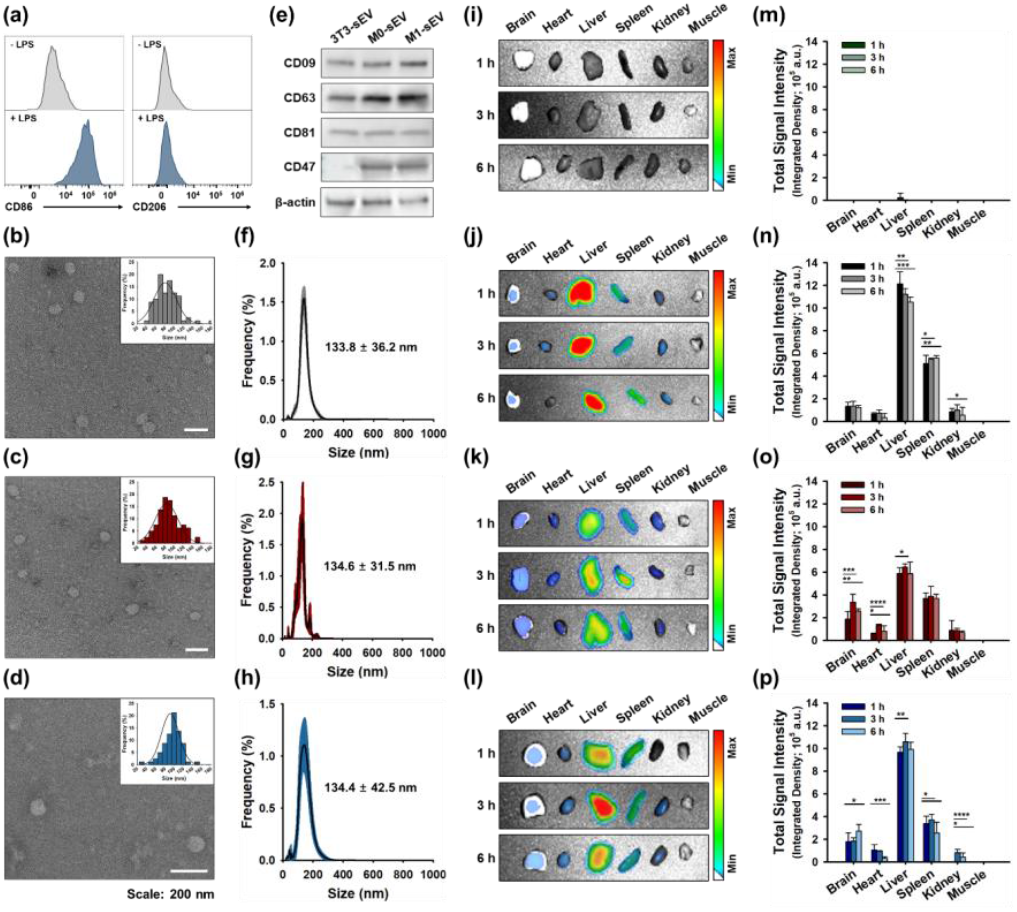
Comprehensive sEV Characterization and M1-sEV Immune Evasion. (a) Fluorescence-activated cell sorting (FACS) analysis of CD86 (M1 marker) and CD206 (M2 marker) in lipopolysaccharide (LPS; 500 ng/mL)-stimulated M0 macrophages to confirm M1 polarization prior to sEV isolation. Transmission electron microscopy (TEM) images showing spherical nanostructures and corresponding size distribution analysis (n = 80) of (b) 3T3-sEVs, (c) M0-sEVs, and (d) M1-sEVs. (e) Western blot analysis of sEV markers. Nanoparticle tracking analysis (NTA) showing size distribution and particle concentration profiles (n = 5) of (f) 3T3-sEVs, (g) M0-sEVs, and (h) M1-sEVs. Representative *ex vivo* fluorescence images and corresponding biodistribution quantification (n = 3, mean ± s.d.) of major organs resected from mice administered with (i, m) Ctrl (PBS), (j, n) 3T3-sEVs, (k, o) M0-sEVs, and (l, p) M1-sEVs. DiD-labeled sEVs (measurement: ex/em = 605/670 nm). Quantification normalized to the fluorescence signal intensity of DiD-3T3-sEVs. One-way ANOVA with Tukey’s post-test. ns: not significant; *p < 0.05; **p < 0.01; ***p < 0.001; ****p < 0.0001.

To further validate the functional role of CD47 in evading immune surveillance during systemic circulation, we performed *ex vivo* fluorescence imaging of major organs (brain, heart, liver, spleen, kidney) and muscle at multiple time points (1, 3, and 6 hours) following tail-vein injection of DiD-labeled sEVs. In the red fluorescence channel, minimal signals were observed in the Ctrl group (Figure 2i, m), which primarily exhibited near-blue background autofluorescence (Figure S3b, c). In the liver, macrophage-derived sEVs showed lower liver retention overall, while 3T3-sEVs exhibited the highest accumulation among all groups (Figure 2j-l). Conversely, spleen accumulation displayed a different trend, with M1-sEVs showing the lowest and 3T3-sEVs the highest levels (Figure 2n-p). These organ-specific distributions can be interpreted within the framework of reticuloendothelial system (RES) clearance. The liver, a central site of innate immune surveillance and cytokine production, (30–32) together with the spleen, which mediates adaptive immunity and antigen presentation, (33–36) exhibited the highest accumulation of 3T3-sEVs, consistent with their rapid opsonization and clearance by immune cells. In contrast, macrophage-derived sEVs (M0- and M1-sEVs) displayed reduced accumulation in both organs, likely attributable to surface CD47 expression that facilitates evasion of immune recognition and suppression of phagocytic clearance. Such RES avoidance underscores the potential of macrophage-derived sEVs as effective carriers for therapeutic delivery. Altogether, these biodistribution findings highlight the stability and immune-evasive properties of macrophage-derived sEVs.

### 3.2. Proteomic Indication for Ligand-Mediated Tumor Homing of M1-sEVs

It has been reported that one of the intrinsic properties of macrophages is their innate immunity-guided tumor-homing behavior, characterized by preferential and site-specific accumulation at tumor sites.(37–39) To investigate whether these properties are also transferred to their derived sEVs, we performed MS-based proteomic analysis to examine their protein composition (Figure 3a). Based on the acquired proteomic profile, we classified several proteins accessions into 13 distinct proteins that potentially reflect tumor specific localization mechanisms: Vinculin, Integrin beta 2 (β2 Integrin), Galectin-3 (Gal-3), Vascular endothelial growth factor receptor 1 (VEGFR1), CD44, Neuropilin-1 (NRP1), Neuropilin-2 (NRP2), Nucleolin, Heat shock protein 70 (HSP70) and 90 (HSP90), Clathrin heavy chain (CHC), Clathrin light chain (CLC), and tyrosine-containing proteins. Among all, Vinculin, stably expressed in most cell types, is widely used as a housekeeping loading control for normalizing protein levels across samples,(40) while both HSP70 and HSP90 are commonly present in sEVs and serve as established markers.(24)

**Figure 3.**
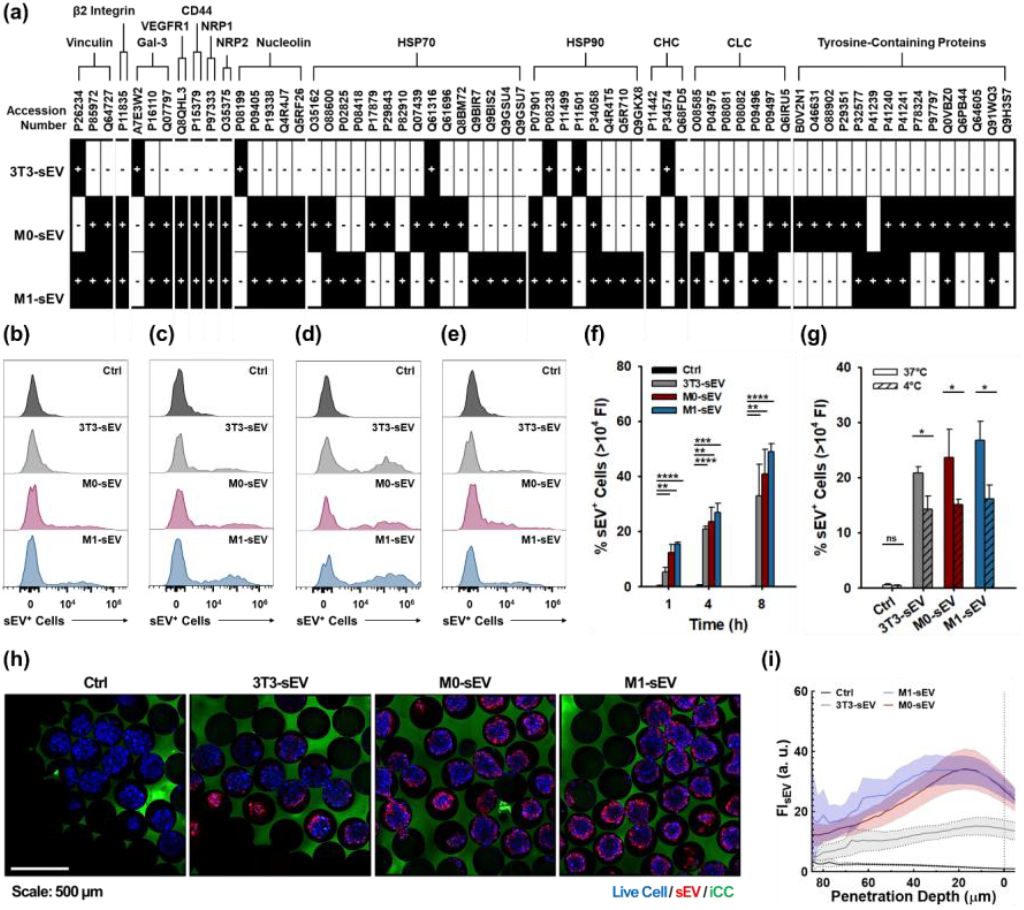
Tumor-homing properties of M1-sEVs. (a) Mass spectrometry (MS)-based proteomic profiling of sEVs. Protein accession numbers from UniProtKB/Swiss-Prot; black boxes with [+]: detected proteins; white boxes with [−]: undetected protein; Gal-3: Galectin-3; VEGFR1: Vascular endothelial growth factor receptor-1; NRP: Neuropilin; HSP: Heat shock protein; CHC: Clathrin heavy chain; CLC: Clathrin light chain. Fluorescence-activated cell sorting (FACS) analysis of DiD-labeled sEV uptake to MCF-7 cells at (b) 1 h, (c) 4 h, and (d) 8 h at 37 °C, with (f) corresponding quantification (n = 3, mean ± s.d.). (e) FACS analysis of temperature-dependent uptake at 4 °C, with (g) corresponding quantification (n = 3, mean ± s.d.). Quantification threshold: DiD fluorescence intensity >10^4^. (h) Two-photon confocal laser scanning microscopy (TP-CLSM) images of DiD-labeled sEV uptake in MCF-7 spheroids in inverted colloidal crystal (iCC) framework (Blue: live cells; Green: 5-DTAF-labeled iCC; Red: DiD-labeled sEV). (i) Quantitatively collected DiD-labeled sEV intensity in the spheroid penetration depth in TP-CLSM images (N > 30). One-way ANOVA with Tukey’s post-test. ns: not significant; *p < 0.05; **p < 0.01; ***p < 0.001; ****p < 0.0001.

More importantly, several expressed proteins on macrophage derived sEVs exhibit functions in directing them toward tumors by mediating interactions with the extracellular matrix (ECM) and components of TME. β2 integrin is a key mediator of leukocyte adhesion, enabling targeted interactions with ICAM-1 and VCAM-1 abundantly present on stromal and endothelial cells in the TME.(41,42) Galectin-3 binds ECM glycoproteins such as laminin, fibronectin, and collagen, facilitating haptotaxis and enhancing their migration and targeting within the TME.(43) VEGF enables responsiveness to VEGF gradients, thereby promoting sEV homing to VEGF-rich TME.(44) CD44 binds specifically to hyaluronic acid (HA) enriched in the tumor ECM, thereby facilitating targeted accumulation within the tumor microenvironment.(45–47) NRP facilitates TME targeting in response to cues such as Semaphorin 3A,(48,49) while NRP1 also serves as a co-receptor for Epidermal growth factor receptor (EGFR) overexpressed in breast, lung, glioblastoma, and colorectal cancers,(50) interacting directly via their extracellular domains.(51,52) In addition, nucleolin was identified, which has been reported to engage in interactions that further target EGFR, particularly through ErbB2.(8,53) The co-expression of NRP1 and nucleolin on macrophage-derived sEVs might enhance their targeted navigation toward EGFR-rich tumor cells and further synergistically act with other ligands involved in ECM interactions and TME recruitment described above.

Another group of proteins potentially enhance uptake at cancer sites through clathrin-mediated endocytosis. CHC forms the structural lattice of the clathrin coat, while CLC regulates clathrin assembly and function.(54) The abundance of these clathrin chains may mimic natural vesicle coats and recruit adaptor proteins such as AP-2,(55) which is enriched at the plasma membrane and initiates clathrin-coated pit formation to drive internalization.(56) In addition, we detected tyrosine-containing proteins, whose tyrosine-based motifs are specifically recognized by AP-2, further facilitating pit formation and cargo uptake.(57) Given that macrophage-derived sEVs present EGFR-targeting ligands, they can engage EGFR to trigger receptor-mediated endocytosis.(8) Thus, the combined presence of clathrin, tyrosine-based sorting signals, and EGFR ligands synergistically promotes clathrin-coated pit formation, enhancing internalization and cellular interactions of macrophage-derived sEVs.

### 3.3 Enhanced Cellular Uptake and Deep Tumor Spheroid Penetration of M1-sEV

Guided by the proteomics results, we assessed whether M1-sEV uptake is enhanced in EGFR-overexpressing cells. To this end, we used MCF-7, a human breast cancer cell line with high EGFR expression,(8) to evaluate internalization *in vitro*. To investigate the kinetics of sEV internalization, MCF-7 cells in 2D culture were incubated with DiD-labeled sEVs for 1, 4, and 8 hours, followed by FACS analysis (Figure 3b–d, f). The percentage of DiD-positive cells increased in a time-dependent manner for all sEV groups. The lowest uptake was observed with 3T3-sEVs (5.30 ± 1.50% at 1 h, 20.83 ± 1.14% at 4 h, and 32.97 ± 11.46% at 8 h), consistent with their minimal expression of EGFR-targeting ligands on the cell surface.(24) Notably, M1-sEVs consistently exhibited the highest uptake at each time point (15.47 ± 0.67% at 1 h, 26.80 ± 3.47% at 4 h, and 48.97 ± 2.90% at 8 h), whereas M0-sEVs showed the second-highest uptake (12.30 ± 2.95% at 1 h, 23.60 ± 5.16% at 4 h, and 40.90 ± 9.01% at 8 h). M1 macrophages, upon activation (e.g., via LPS stimulation), upregulate EGFR-associated ligands such as heparin-binding EGF-like growth factor (HB-EGF) relative to M0 macrophages.(58,59) These results indicated that M1-sEVs may exhibit higher levels of these ligands compared to the M0-sEVs. To further examine whether sEV uptake involves active transport via endocytosis, MCF-7 cells were incubated with DiD-labeled sEVs at 37 °C or 4 °C for 4 hours (Figure 3e, g). FACS analysis revealed that all sEVs uptake was substantially lower at 4 °C compared with 37 °C, while no significant difference was observed among three groups. These results indicated that the increased uptake of M1-sEVs at 37 °C is driven by an energy-dependent process, consistent with active endocytosis facilitated by EGFR-associated ligand–receptor interactions.

To examine how targeted uptake is associated with ECM and TME recruitment-associated ligands, we investigated sEV penetration into tumor spheroids in 3D cell culture *in vitro*. We conducted a proof-of-concept experiment using MCF-7 spheroids generated in a high-yield iCC framework established previously.(21) After 6 days (144 h) of culture, spheroids were treated for 1 day (24 h) with DiD-labeled sEVs, followed by PFA fixation. TP-CLSM imaging revealed origin-dependent differences in penetration (Figure 3h), with notably stronger red fluorescence signals from macrophage-derived sEVs, particularly M1-sEVs, indicating enhanced uptake accompanied by highly efficient penetration. Quantitative spatial analysis using our previously developed pipeline(22) further confirmed that M1-sEVs penetrated deeper into MCF-7 spheroids than M0-sEVs and 3T3-sEVs (Figure 3i, S5), suggesting a synergistic effect of tumor-homing proteins on M1-sEVs on enhancing sEV uptake in tumor tissue. Collectively, these results indicated that ligand-mediated tumor homing of M1-sEVs not only enhanced their uptake and promoted strong infiltration into 3D spheroids.

### 3.4. Tumor-suppressive miRNAs and Their Anti-tumor Effects on Breast Cancer

As several tumor-suppressive miRNAs have been reported in M1 macrophages and their sEVs,(15,16,60,61) which modulate cancer progression, we investigated their collective functional effects on breast cancer. To explore the miRNA cargos of sEVs, we first confirmed the quality of the extracted RNA using automated electrophoresis with High Sensitivity RNA ScreenTape (Figure S6b-e) and quantified the amount of extracted RNA from each sEV. qRT-PCR was then performed to quantify miRNAs associated with cancer suppression, including those reported to express in M1 macrophages.(15,16,61) To normalize RNA input, we first confirmed consistent U6 levels across all sEVs (Figure S6a). miR-150 expression in M1-sEVs was then examined for both mature miRNA molecules derived from the same pre-miRNA hairpin but originating from opposite arms (5′ and 3′). The levels of miR-150-5p (Figure 4a) were similar between 3T3-sEVs and M0-sEVs, but were significantly elevated in M1-sEVs, showing 9.96 ± 3.12-fold higher compared to that of 3T3-sEVs. For miR-150-3p (Figure 4b), M0-sEVs showed 2.20 ± 0.21-fold higher than 3T3-sEVs, while M1-sEVs exhibited a 263.65 ± 20.35-fold higher abundance. miR-150 is known to suppress tumor progression by inhibiting proliferation, migration, and invasion through targeting MMP16,(16) which is upregulated in breast cancer cells and promotes metastasis.(62) Similarly, miR-181a-5p was highly enriched in M1-sEVs (Figure 4c), showing a 36.59 ± 1.79-fold increase compared to 3T3-sEVs, with M0-sEVs showing only 2.25 ± 1.01-fold increase. miR-181a has been reported to influence cancer cells via the STK16 pathway, thereby regulating cell viability and apoptosis.(15) Notably, MCF-7 cells are highly responsive to STK16-mediated mechanisms, which control cell number and the accumulation of binucleated cells,(63) potentially leading to cancer growth-inhibitory effects. Moreover, miR-34 levels were significantly elevated in M1-sEVs.(61) For miR-34a-5p (Figure 4d), M1-sEVs exhibited a 16.89 ± 0.90-fold increase compared to 3T3-sEVs, while M0-sEVs showed a 3.44 ± 0.19-fold increase. Similarly, for miR-34a-3p (Figure 4e), M1-sEVs expressed 27.71 ± 4.33-fold higher levels than 3T3-sEVs, whereas M0-sEVs displayed only a 1.76 ± 0.31-fold increase. It is well established that miR-34 functions as a tumor suppressor by regulating epithelial–mesenchymal transition (EMT) through EMT-associated transcription factors and p53, key drivers of tumor progression, metastasis, and therapy resistance.(64) Collectively, we identified miRNAs passed down from M1 macrophages into M1-sEVs, which the strongest anticancer activity among all sEV types. Consistently, TP-CLSM imaging (Figure S7) showed greater shrinkage of tumor spheroids in the M1-sEV group (Figure 4f), confirming that M1-sEVs possess both tumor-homing ability and growth-inhibitory effects, enabling enhanced tumor suppression in 3D breast cancer models *in vitro*.

**Figure 4.**
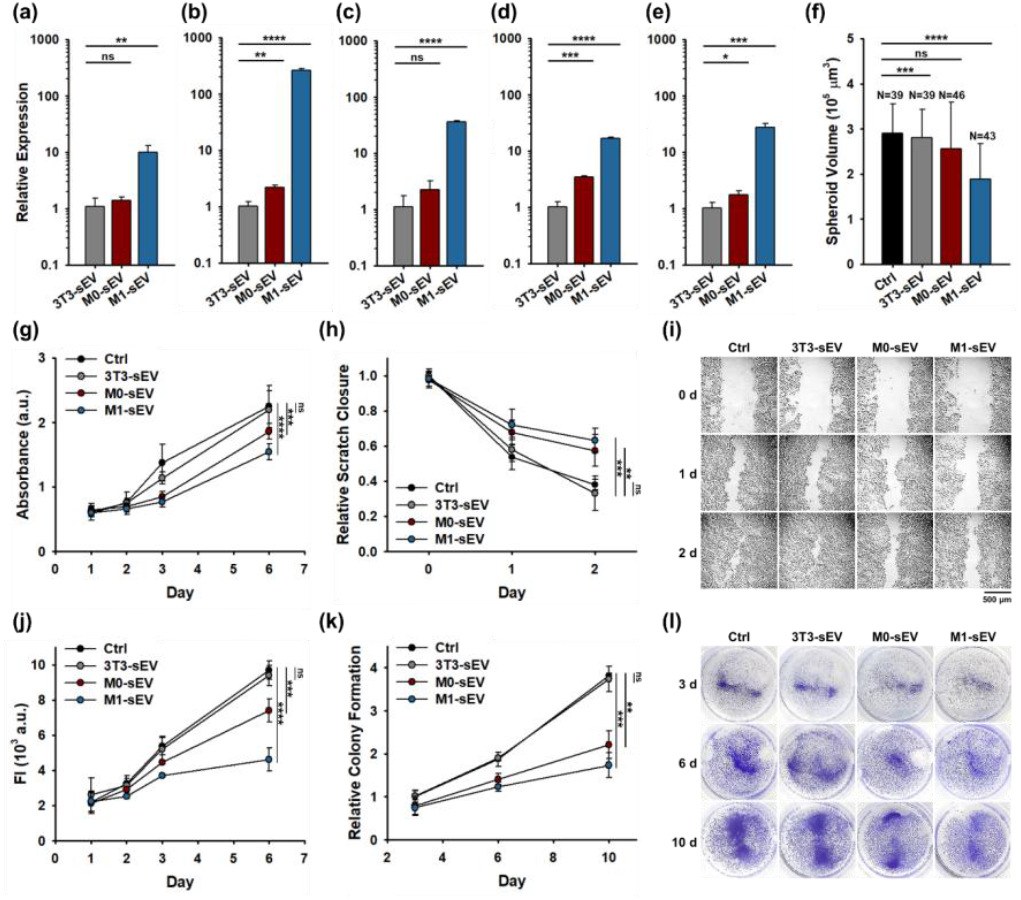
Anti-cancer miRNAs in M1-sEVs and their functional effects. Quantitative reverse transcription polymerase chain reaction (qRT-PCR) analysis of (a) mmu-miR-150-5p, (b) mmu-miR-150-3p, (c) mmu-miR-181a-5p, (d) mmu-miR-34a-5p, and (e) mmu-miR-34a-3p expression levels in sEVs (n = 3, mean ± s.d.). Comparable U6 levels across the three sEV groups were confirmed as housekeeping gene. Gene expression is normalized to miRNA levels in 3T3-sEVs and shown as relative changes. (f) Quantification of spheroid volume from TP-CLSM images. Evaluation of anti-proliferative effects via (g) Water-Soluble Tetrazolium 8 (WST-8) assay (n = 4, mean ± s.d.), (h, i) scratch assay (n = 4, mean ± s.d.), (j) resazurin assay (n = 4, mean ± s.d.), and (k,l) clonogenic assay (n = 3, mean ± s.d.). One-way ANOVA with Tukey’s post-test. ns: not significant; *p < 0.05; **p < 0.01; ***p < 0.001; ****p < 0.0001.

To further verify whether the miRNAs in M1-sEVs affect breast cancer proliferation, *in vitro* functional assays, WST-8 and resazurin assays, were performed in MCF-7 cells. The WST-8 assay measures cell viability based on mitochondrial dehydrogenase activity, reflecting the overall metabolic activity.(65) In contrast, the resazurin assay detects the reduction of resazurin to resorufin by a broader range of cellular metabolic processes, making it more sensitive to subtle changes in proliferation and metabolic stress.(66) In the WST-8 assay (Figure 4g), treatment of MCF-7 cells with sEVs showed no significant differences compared to the non-treated control group until day 2. However, from day 3 onward, macrophage-derived sEVs began to diverge from both the control and 3T3-sEV groups. By day 6, M1-sEVs resulted in a lower cell viability compared to M0- and 3T3-sEVs (Figure S8a). Correspondingly, the resazurin assay (Figure 4j) reflected the same trend as the WST-8 results. The relative fluorescence intensity increased on day 6 by 4.44 ± 0.24-fold in the control group, 4.31 ± 0.26-fold in 3T3-sEV, 3.39 ± 0.15-fold in M0-sEV, and, notably, only 2.12 ± 0.30-fold in M1-sEV-treated cells (Figure S8b). The inhibitory effect observed in the functional assays demonstrated the cumulative impact of tumor-suppressive miRNAs, including miR-150, miR-181a, and miR-34a, delivered by M1-sEVs.

To assess the consequent inhibitory effects on cancer progression in MCF-7 cells, we performed a scratch assay (Figure 4h, i) to evaluate cell adherence and migration, indicative of proliferative and metastatic potential. Control and 3T3-sEV treated MCF-7 cells exhibited rapid wound closure, with comparable migration rates (61.98 ± 3.03% and 66.92 ± 9.94%, respectively). M0-sEV treatment moderately impaired closure, with 42.48 ± 8.98% closure at the same time point. M1-sEV treatment showed the most pronounced inhibition of scratch closure, with only 36.76 ± 6.95% closure at Day 2, indicating strong suppression of migratory activity (Figure S8c). These results suggested that M1-sEVs exert the greatest inhibitory effect on cell motility, thereby potentially limiting metastatic progression. Moreover, we performed a clonogenic assay to assess the long-term proliferative potential of MCF-7 cells following sEV treatment.(8) MCF-7 cells were treated with sEVs for 3 days, then reseeded at equal density, and colonies were observed at 3, 6, and 10 days post-reseeding (Figure 4k, l). Control and 3T3-sEV treated groups formed numerous colonies, with fold changes of 3.82 ± 0.05 and 3.73 ± 0.29, respectively, indicating robust proliferative capacity. M0-sEV treatment somewhat reduced colony formation (2.21 ± 0.32-fold change), whereas M1-sEV treatment showed the most pronounced inhibition, with only 1.73 ± 0.29-fold change in colony number (Figure S8d). As a reduction in colony number or size reflects decreased proliferative potential or survival, these results suggested that M1-sEVs strongly suppress the long-term growth capacity of breast cancer cells.

### 3.5. Synergistic Anticancer Effects of Dox loaded M1-sEVs

To load Dox into sEVs, we utilized the established method using chiral GQDs, as reported in our previous studies (see SI Methods and Results, Figures S9–S14).(19,67) The loading efficiency of Dox was determined to be 62.53 ± 11.85% for 3T3-sEVs, 63.94 ± 5.12% for M0-sEVs, and 63.36 ± 2.65% for M1-sEVs (Figure 5a, b), which significantly surpass the common sEV loading methods.(7,19) To explore the dual function of M1-sEVs with their inherent antitumor activity and drug delivery, we evaluated the *in vitro* cytotoxicity of Dox-loaded sEVs (Dox/sEV) in MCF-7 cells (Figure S15d). At 24 h (Figure 5c, S16a-b), the half maximal inhibitory concentration (IC_50_) of free Dox was 2.10 µM, whereas Dox/M1 sEVs showed a lower IC_50_ of 1.45 µM, followed by Dox/M0 sEVs at 1.54 µM and Dox/3T3 sEVs at 1.89 µM. After 48 h (Figure 5d, S16c), the differences became more pronounced with free Dox at 1.45 µM compared with 0.46 µM for Dox/M1 sEVs, 1.35 µM for Dox/M0 sEVs and 1.42 µM for Dox/3T3 sEVs. Since Dox inhibits cell proliferation by intercalating into nuclear DNA and blocking replication and transcription, longer exposure allowed Dox to more effectively suppress essential gene activity.(8) This effect was further enhanced by intrinsic anti-tumor effect of M1-sEVs, resulting in greater Dox sensitivity at 48 h compared with 24 h.

**Figure 5.**
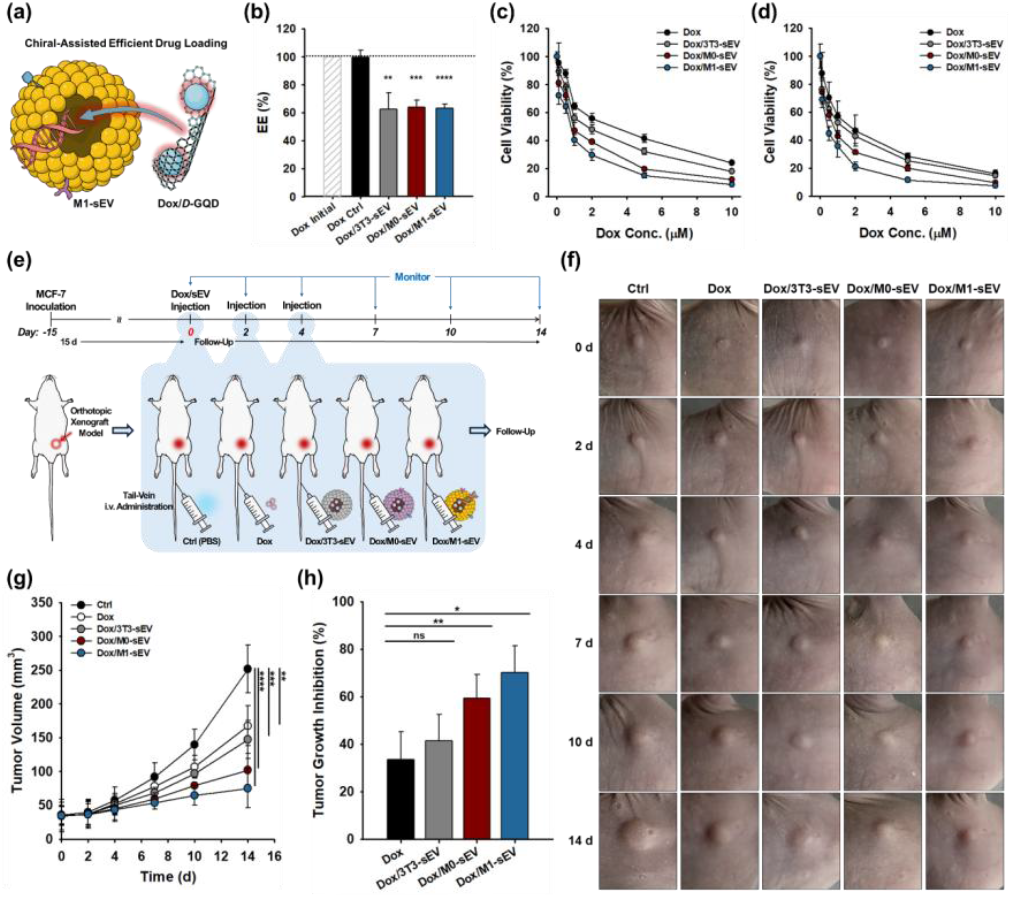
Synergistic therapeutic efficacy via the combination of M1-sEV–mediated intrinsic anticancer potential and their capacity for drug delivery. (a) Schematic illustration of Doxorubicin (Dox) loading into sEVs using chiral graphene quantum dots (GQDs). (b) Quantification of Dox encapsulation efficiency (EE%). Cell viability of MCF-7 cells treated with Dox-loaded sEVs at various concentrations (0–10 μM) for (c) 24 h and (d) 48 h (n = 4, mean ± s.d.). (e) Schematic workflow for *in vivo* therapeutic efficacy evaluation of Dox-loaded sEVs. (f) Representative photographic images of MCF-7 tumor-bearing female NCr nude mice taken at 0, 2, 4, 7, 10, and 14 days post-injection. (g) Corresponding tumor volume monitoring and (h) tumor growth inhibition rate (TGI%) at day 14 (n = 4, mean ± s.d.).Intravenous (i.v.) injection of Dox, 4 mg/kg; administered every 2 days; total of three doses; Dox concentration in sEVs determined by encapsulation efficiency (EE%). One-way ANOVA with Tukey’s post-test. ns: not significant; *p < 0.05; **p < 0.01; ***p < 0.001; ****p < 0.0001.

Lastly, we validated the feasibility of the combinational Dox/M1-sEV system *in vivo* using MCF-7 tumor-bearing orthotopic xenograft mouse models. Mice received three tail-vein intravenous injections of either control (PBS), free Dox, or Dox loaded into three types of sEVs (4 mg/kg Dox per dose). Tumor progression was monitored for 2 weeks following the first administration (Figure 5e). As shown in Figures 5f and g, the Dox/M1-sEV group exhibited the most pronounced tumor growth suppression, followed by Dox/M0-sEV, Dox/3T3-sEV, and free Dox, consistent with the *in vitro* cytotoxicity results (Figure S17). The tumor growth inhibition rates (TGI%) were 33.48 ± 11.73 for Dox, 41.50 ± 11.07 for Dox/3T3-sEV, 59.45 ± 9.84 for Dox/M0-sEV, and 70.18 ± 11.37 for Dox/M1-sEV (Figure 5h), highlighting the superior therapeutic efficacy of the Dox/M1-sEV treatment. Furthermore, histological analysis of major organs (heart, lung, liver, spleen, and kidney) and muscle revealed no visible damage in any treatment group (Figure S18a). In contrast, tumor tissues exhibited significant differences among groups (Figure S18b). The control group showed intact tumor cell nuclei and well-organized tissue architecture. The free Dox group displayed evident nucleus damage and tissue disorganization, with a swirled morphology. The Dox/3T3-sEV group presented a similar pattern to free Dox. The Dox/M0-sEV group exhibited more pronounced tissue deterioration and a substantial reduction in nuclear density, likely due to enhanced tumor cell uptake, as M0-sEVs express tumor-homing ligands toward MCF-7 cells. Notably, the Dox/M1-sEV group showed the most severe tumor damage, characterized by fragmented, splintered, and scattered nuclei, which were attributed to the synergistic effects of anti-tumor miRNAs carried by M1-sEVs along with tumor tumor-homing effect. These findings aligned with the tumor photographs (Figure 5f), showing the most pronounced tumor growth suppression and visibly darker tumor tissue in the Dox/M1-sEV group. Notably, tumors in M1-sEV– treated mice appeared reddish-brown (Figure 5f), likely reflecting necrosis induced by their strong anticancer effects, which often appears dark due to blood accumulation or hemoglobin breakdown.(68,69) These results further align with photographic and histological analyses, where tumors from the Dox/M1-sEV group (Figure S18b) exhibited nuclear deformation, disorganized architecture, and amorphous eosinophilic debris, classic hallmarks of severe necrosis that were more pronounced than in other groups. Taken together, this outcome demonstrates the effectiveness of dual role M1 sEV drug delivery system in harnessing their innate biological advantages, particularly intrinsic anticancer activity. At the same time, it also made it possible to fully exploit the drug encapsulation and delivery capabilities of sEVs via the chiral-assisted drug loading technique, thereby achieving the most significant therapeutic effect in MCF-7 tumor-bearing mice.

## 4. DISCUSSION

In this study, we investigated the dual function of M1-sEVs as innate anticancer agents and targeted drug delivery nanocarriers to achieve a synergistic therapeutic effect. Retaining the key functions of their parental M1 macrophages, we demonstrated that M1-sEVs act as highly efficient dual therapeutic nanoplatforms that integrate their intrinsic bioactivity with chemotherapeutic drug delivery, thereby offering distinct advantages over conventional pharmacological approaches.

The first advantage we demonstrated is that M1-sEVs leverage autologous membrane proteins and lipid composition inherited from their parental M1 macrophages, allowing them to evade immune surveillance, and resist clearance by the mononuclear phagocyte system (Figure 2l, p). In contrast, free drugs face rapid clearance, poor stability, and limited tumor accumulation,(70) while synthetic nanoparticles, such as lipid nanoparticles (LNPs), improve drug delivery, but are often rapidly recognized by the reticuloendothelial system (RES), leading to off-target accumulation in the liver and spleen.(71) Here, we demonstrated that M1-sEVs exhibited superior pharmacokinetics and biodistribution, particularly compared to sEV derived from non-immune cells (3T3-sEVs; Figure 2i, n). Building on their immune cell origin, M1-sEVs inherit tumor-targeting ligands, such as NRP1 and nucleolin (Figure 3a) from their parent cells, enabling preferential homing to EGFR-expressing tumors. This effect is further enhanced by additional endogenous ligands, including β2 integrin, Gal-3, VEGFR1, CD44, and NRP1 (Figure 3a), which engage TME components such as ICAM-1, VCAM-1, glycans, VEGF, HA, and Semaphorin 3A, collectively guiding M1-sEVs migration and promoting tumor-specific homing. Considering that the extracellular components of the TME are genetically more stable than those expressed by very heterogeneous tumors, sEVs leverage this inherent complementarity to achieve superior tumor tropism through sustained engagement.(72) In contrast, LNPs are typically engineered with one or two ligands for single-pathway targeting.(73) Even though multi-ligand LNPs have been reported, they often face rapid opsonization, Kupffer cell uptake, and liver accumulation, as their exogenously added decorations remain vulnerable to immune surveillance.(74,75) Hence, the innate multivalent recognition of sEVs through targeting ligands, multiple integrins, and adhesion molecules enables simultaneous engagement with tumor cells, TME components, and ECM proteins, creating redundancy that strengthens binding and ensures robust tumor homing.(76)

We further investigated the antitumor properties of M1-sEVs inherited from their parental M1 macrophages. In particular, we identified a set of miRNAs highly enriched in M1-sEVs, sorted from their cellular origin, that likely contribute to their antitumor effects. Compared with 3T3-sEVs and M0-sEVs, M1-sEVs were markedly enriched in multiple tumor-suppressive miRNAs, including miR-150-5p (9.96 ± 3.12-fold), miR-150-3p (263.65 ± 20.35-fold), miR-181a-5p (36.59 ± 1.79-fold), and miR-34a-5p/3p (16.89 ± 0.90- and 27.71 ± 4.33-fold; Figures 4a–e). These miRNAs regulate proliferation, migration, invasion, and EMT through distinct but complementary mechanisms: miR-150 via MMP16,(16) miR-181a via STK16,(17) and miR-34a via p53/EMT,(61) all of which are detected at very high levels in M1-sEVs. In addition, miR-628, also reported to be retained in M1-sEVs, acts via METTL14/m6A signaling in ERα-positive breast cancer cells.(15) Collectively, their downstream targets converge on critical oncogenic pathways, functioning in parallel and intersecting networks to inhibit tumor progression. On top of that, the natural enrichment of anticancer miRNAs in M1-sEVs, co-encapsulated with multiple tumor-suppressive genes, provides an inherent advantage for streamlined nanotherapeutic design. In contrast, synthetic nanocarriers, such as polymersome and liposome, typically require complex chemical and physical strategies to achieve efficient co-loading while protecting genomic cargos from degradation or denaturation.(77,78) Thus, while leveraging their endogenous enrichment of synergistic anticancer miRNAs, M1-sEVs are able to provide reproducible, bioinert cargo delivery establishing them as an efficient and safe platform for combinatorial gene therapy.

sEVs inherently protect therapeutic cargo from enzymatic degradation, enhance bioavailability through targeted uptake, and are biodegradable, avoiding the accumulation issues seen with synthetic nanoparticles.(79) Their delivery efficiency is further supported by multiple uptake pathways and partial lysosomal escape, which can be further enhanced by surface engineering.(8,80) Leveraging these advantages, we successfully loaded doxorubicin (63.36%) into M1-sEVs using our chiral GQD-based technology (Figure 5a, b),(8,19) achieving high encapsulation while preserving sEV integrity and bioactivity. This dual-function system—combining intrinsic anticancer activity with targeted drug delivery—produced a 3-fold reduction in IC_50_ *in vitro* (0.46 μM vs. 1.45 μM; Figure 5d) and 70.18% tumor growth inhibition *in vivo*. These findings highlight the potent anticancer efficacy of M1-sEVs and demonstrate their promise as an advanced nanotherapeutic platform that integrates endogenous bioactivity with chemotherapeutic delivery.

Lastly, M1-derived sEVs have been reported to deliver therapeutic cargos to otherwise difficult-to-reach tissues, including penetration of dense tumor stroma and traversal of the blood–brain barrier (BBB).(81) Consistent with these, our *ex vivo* biodistribution (Figure 2i–p) demonstrated significantly higher brain accumulation of macrophage-sEVs compared to 3T3-sEVs. M1-sEVs express EGFR-targeted ligands, which are also overexpressed in glioblastoma,(51) and carry miR-150 that suppresses glioma progression (Figure 4a, b). These features highlight their strong potential for expanding M1-sEV–based therapeutic platforms in neurological cancer treatment.

## 5. CONCLUSION

In this study, we collectively demonstrated that M1-sEVs not only retained the tumor-suppressive bioactive properties of their parent cells but also served as efficient drug carriers, providing multifunctional anticancer therapy. M1-sEVs exhibited high stability in circulation due to their immune surveillance escape, and intrinsic tumor-homing capabilities, enabling efficient uptake by breast cancer cells and deep penetration into tumor spheroids. In addition, they carried anti-proliferative miRNAs that collectively suppressed cancer cell growth, metabolic activity, and key processes associated with tumor progression, including migration, invasion, adhesion, and self-renewal. Through this convergence of their inherent anticancer effect and drug payload capability, dual role M1-sEVs significantly enhanced the anticancer efficacy in both *in vitro* with ∼3-fold reduction in IC_50_ and achieved a 70.18% TGI *in vivo* relative to control. Overall, this study established M1-sEVs as dual therapeutic platforms that combined innate tumor-suppressive activity with enhanced chemotherapeutic delivery, providing a versatile foundation for cancer therapy and next-generation combination approaches.

## AUTHOR INFORMATION

### Authors

Gaeun Kim - Department of Chemical and Biomolecular Engineering, University of Notre Dame, Notre Dame, Indiana 46556, United States; Harper Cancer Research Institute, University of Notre Dame, Notre Dame, IN 46556, United States;

Hyunsu Jeon - Department of Chemical and Biomolecular Engineering, University of Notre Dame, Notre Dame, Indiana 46556, United States;

Adrian Chao - Department of Biological Sciences, University of Notre Dame, Notre Dame, Indiana 46556, United States

James Johnston - Department of Chemical and Biomolecular Engineering, University of Notre Dame, Notre Dame, Indiana 46556, United State

Runyao Zhu - Department of Chemical and Biomolecular Engineering, University of Notre Dame, Notre Dame, Indiana 46556, United States; orcid.org/0009-0005-5109- 0020

Courtney Khong - Department of Chemical and Biomolecular Engineering, University of Notre Dame, Notre Dame, Indiana 46556, United States

Yichen Liu - Department of Chemical and Biomolecular Engineering, University of Notre Dame, Notre Dame, Indiana 46556, United States

Minzhi Liang - Department of Biological Sciences, University of Notre Dame, Notre Dame, Indiana 46556, United States

Xin Lu - Department of Biological Sciences, University of Notre Dame, Notre Dame, Indiana 46556, United States; Harper Cancer Research Institute, University of Notre Dame, Notre Dame, IN 46556, United States

### Author Contributions

YW and GK conceptualized the study. GK designed the methodology and carried out all the experiments, as well as all data analysis. HJ conducted and analyzed 3D spheroid experiments. AC conducted qPCR for the identification of miRNA in sEVs under the supervision of XL. JJ isolated M1-sEVs and performed ELISA analysis. RZ synthesized and characterized chiral GQDs, and contributed to the *ex vivo* study. CK took the TEM images. YL assisted *in vivo* study. ML provided technical support for animal study under the supervision of XL. GK wrote the original manuscript and created all schematic illustrations in this manuscript. YW supervised the entire project, including manuscript writing. All authors contributed to data interpretation, discussion, and writing.

## Acknowledgements

TEM, CLSM, *ex vivo*, and H&E histology images were obtained using the instruments of Notre Dame Integrated Imaging Facility (NDIIF), University of Notre Dame. NTA analysis was obtained using the instrument of Harper Cancer Research Institute (HCRI), University of Notre Dame. MS analysis for sEV proteins was conducted by the instrument of Mass Spectrometry & Proteomics Facility (MSPF), University of Notre Dame. FTIR analysis was obtained using the instrument at the Center for Environmental Science and Technology (CEST), University of Notre Dame. Western blot images were acquired using the instrument of Biophysics Instrumentation Core (BIC), University of Notre Dame. RNA QC was conducted by Genomics & Bioinformatics Core Facility (GBCF), University of Notre Dame. All animal studies were conducted at Freimann Life Science Center (FLSC), University of Notre Dame. We appreciate the support from these facilities for this study. This study was supported in part by the Berry Family Foundation Graduate Fellowship from the Berthiaume Institute for Precision Health (BIPH; to GK) at the University of Notre Dame. This study was supported in part by the Interdisciplinary Interface Training Program (IITP) Grant from HCRI, with funding from the Walther Cancer Foundation (WCF; to GK).

## Statements and Declarations

### Ethical Approval Statement

Animals involved in this study under protocol 22-08-7344 and 24-11-8960 that reviewed and approved by The University of Notre Dame’s Institutional Animal Care and Use Committee (IACUC).

## Table of Contents

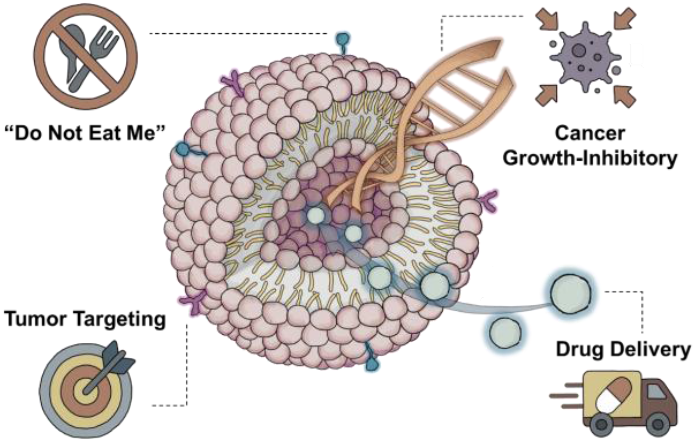

M1 macrophage-derived small extracellular vesicles (M1-sEVs) exhibit intrinsic antitumor activity by evading immune surveillance, homing to tumors, and suppressing cancer growth. Leveraging these functions alongside their capacity to deliver therapeutic payloads, dual-action M1-sEVs achieve enhanced efficacy in vitro and in vivo. This study establishes M1-sEVs as a versatile nanoplatform, combining innate tumor suppression with chemotherapeutic delivery for next-generation combination cancer therapy.

