## Supporting Information for "M1 Macrophage-Derived Small Extracellular Vesicles as Synergistic Nanotherapeutics: Harnessing Intrinsic Anticancer Activity and Drug Delivery Capacity"

### Supplementary Methods

#### Synthesis and Characterization of Chiral Graphene Quantum Dots (GQDs)

The GQDs were synthesized using a modified Hummers' method based on our previous report as follows:[1,2] Briefly, carbon nanofibers were used as a precursor for the top-down synthesis of GQDs via oxidative treatment with $H_2SO_4$ (98%) and $HNO_3$ (68%) at 120 °C for 20 h. The product was further neutralized by NaOH solution and then purified by dialysis using a dialysis bag with a 1 kDa molecular weight cutoff (MWCO). Surface modification with *D*-cysteine was then performed via a 1-ethyl-3-(3-dimethylaminopropyl)carbodiimide (EDC) and N-hydroxysuccinimide (NHS) coupling reaction. The synthesized chiral GQDs were further purified using the same dialysis method and filtered through a 0.22 μm syringe filter. Characterization of the chiral GQDs was conducted as follows: Nanostructure and particle size were assessed using transmission electron microscopy (TEM) at an accelerating voltage of 200 kV. Fluorescence emission was measured with a microplate reader. Optical absorption and chiroptical properties were evaluated using circular dichroism (CD) spectroscopy, and chemical composition was analyzed using Fourier-transform infrared spectroscopy (FTIR) spectroscopy.

### Supplementary Results

#### Additional Validation of LPS-Induced M1 Macrophage–Derived sEVs

Furthermore, we found that both macrophage-derived sEVs contain a greater variety of HSP70 types compared with 3T3-sEVs in MS-based proteomics (**Figure 3a**). The number of HSP70 types was similar between M0-sEVs (8 protein accessions) and M1-sEVs (9 protein accessions), with only 2 overlapping, indicating that HSP70 composition changes following LPS-induced differentiation. As HSP70 has been reported to be a key molecule regulating lipid metabolism in decidual macrophages and influencing their function,[3] the diversity of HSP70s in macrophage-derived sEV likely underlines the mechanism for immune-regulatory properties of macrophage-derived sEVs. It has been reported that Toll-like receptor (TLR) signaling, particularly TLR4–NF-κB, is stimulated by LPS and other microbial ligands.[4] This subsequently activates macrophages, including RAW 264.7 cells which used in our study, driving their polarization toward an M1 phenotype.[5] Given another report showing that HSP90 inhibitors repress TLR4-mediated NF-κB activity,[6] it is likely that macrophages upregulate HSP90 to stabilize key factors in the TLR4–NF-κB pathway, thereby facilitating M1 differentiation in response to LPS.

#### Comprehensive Characterization of Chiral GQDs

Chiral GQDs, a highly efficient platform for Dox loading into sEVs, were developed by modifying GQDs with chiral amino acids, such as cysteine, as reported in our previous studies.[1,7] In the CD spectra, chiral GQDs functionalized with *D*-cysteine (*D*-GQDs) exhibited distinct chiroptical activity (**Figure S9a**), and their fluorescence emission spectrum (**Figure S9b**) was red-shifted relative to unmodified GQDs, likely due to the electron-donating effect of *D*-cysteine. This observation was further supported by the shifted absorbance spectra **(Figure S9c**), confirming the successful chiral functionalization of GQDs. The chemical composition of *D*-GQDs was analyzed using FTIR spectroscopy (**Figure S9d**), showing prominent peaks at 1706 $cm^{-1}$ (C=O stretching) and 1250 $cm^{-1}$ (C–N stretching attributed to *D*-cysteine), compared to GQDs. Quantitative TEM analysis (**Figure S9e**) revealed an average diameter of 6.44 ± 2.54 nm for *D*-GQDs, which was intentionally tuned to match the lipid bilayer thickness for optimal permeability into sEVs and to enable rapid renal clearance, as shown by *ex vivo* biodistribution (**Figure S10**).

**Efficient Doxorubicin (Dox) Loading into sEVs via a Chiral GQD Nanoplatform**

This size-tuned *D*-GQD forms a complex with Dox through pi stacking and has been reported to exhibit high permeation of this complex into sEVs.[1] This is attributed to selective nanoscale interactions between twisted nanosheet-structured *D*-GQDs and the lipid bilayers of sEVs, which, in turn, preserve the structural integrity of sEVs without compromise.[1,8,9] We observed that the fluorescence intensity of Dox decreased at a fixed concentration of 200 μM upon titration with increasing concentrations of *D*-GQDs (0–22.5 μM). This indicated successful complex formation, and the estimated complex ratio suggested that each *D*-GQD could carry approximately 14 Dox molecules (**Figure S11**). The structural integrity of sEVs after Dox loading was confirmed by NTA (**Figure S12a**), showing no significant change in size distribution, and further validated by SMLM (**Figure S12b**), which revealed colocalization of the sEV membrane (DiO; green) with *D*-GQDs (blue) and Dox (red) in individual images. The successful loading of Dox into sEVs via *D*-GQDs was validated by significant fluorescence recovery (**Figure S13**). This was assessed as the change in emission intensity of DiO-stained sEV membranes before and after lysis, alongside a notable increase in Dox fluorescence upon release from sEVs. Both effects are attributed to the reduced self-quenching of DiO and Dox on and in sEVs, respectively, resulting in enhanced fluorescence intensity.[1,8,9]

**Quantification of Dox Encapsulation Efficiency in sEVs**

To quantify the encapsulation efficiency of Dox in sEVs, we modified a method developed for assessing drug complexation efficiency on graphene oxide with chemotherapeutic agents.[10] After loading Dox into sEVs via *D*-GQDs, the lipid membrane of the sEVs was lysed to release the encapsulated cargo. The extracted Dox/*D*-GQD complexes were then dissolved in acetonitrile and sonicated to disrupt pi stacking interactions between Dox and GQDs. Acetonitrile served as a solvent that influences the molecular environment, reducing hydrogen bonding and other interactions, thereby facilitating the disruption of π–π stacking.[11] The detached Dox was separated from *D*-GQDs by filtration through a 2 kDa membrane (**Figure S14a**). The fluorescence of the filtrate-eluent containing free Dox was measured, and Dox concentration was quantified using a calibration curve (**Figure S14b**) of fluorescence intensity versus Dox concentration. Based on this calibration, the loading efficiency of Dox was determined to be 62.53 ± 11.85% for 3T3-sEVs, 63.94 ± 5.12% for M0-sEVs, and 63.36 ± 2.65% for M1-sEVs (**Figure 5**).

**Supplementary Figures**

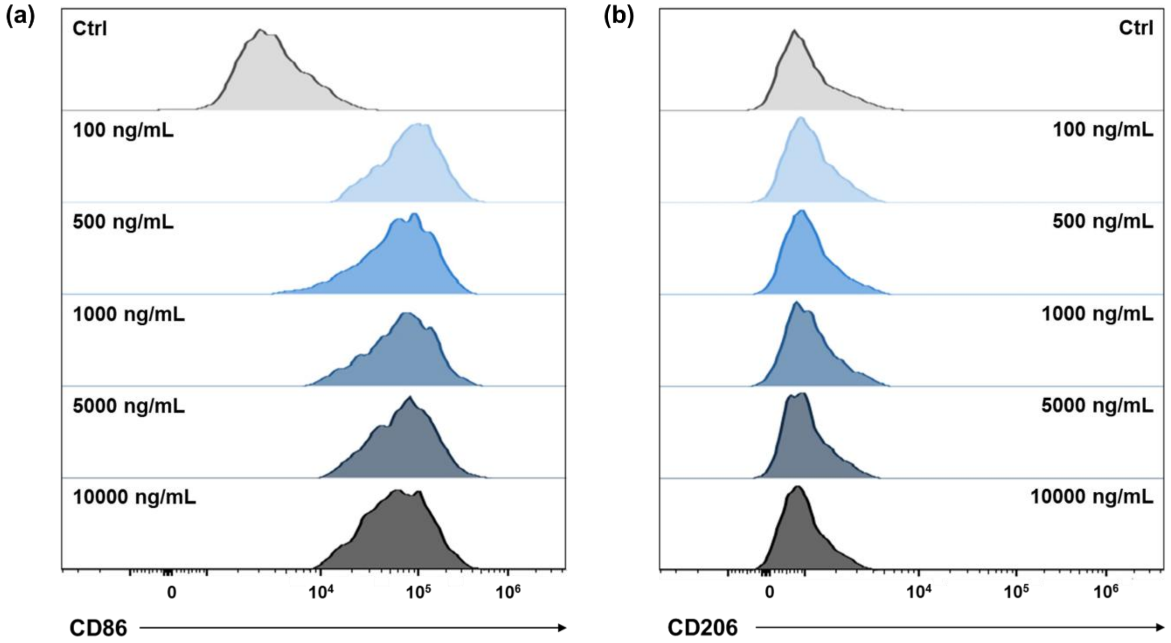


**Figure S1**. Fluorescence-Activated Cell Sorting (FACS) analysis of (a) CD86 (M1 marker) and (b) CD206 (M2 marker) in M0 macrophages treated with varying concentrations of Lipopolysaccharide (LPS) to confirm M1 polarization prior to small extracellular vesicle (sEV) isolation.

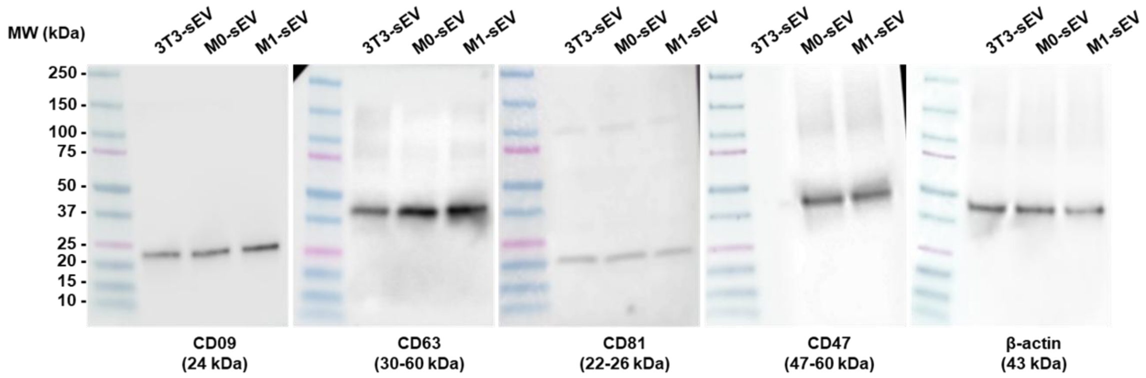


**Figure S2**. Full membrane Western blot of sEV markers.

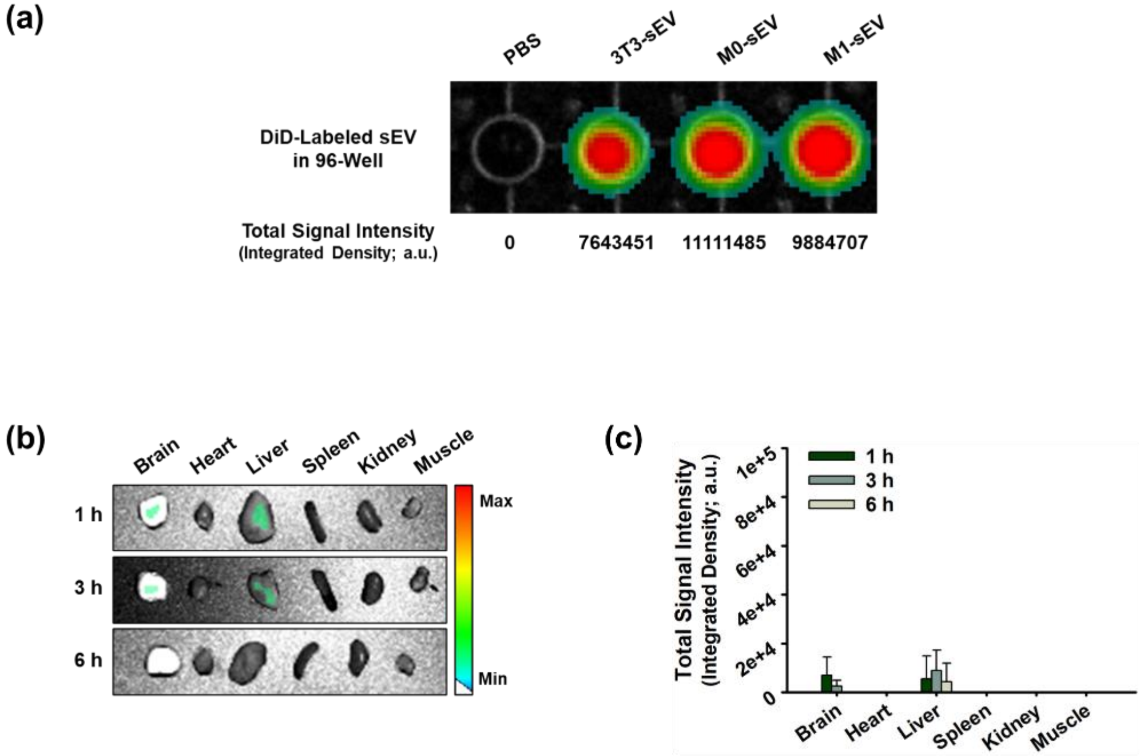


**Figure S3**. (a) Fluorescence intensity of DiD-sEV in 96-well plate for normalization of *ex vivo* biodistribution quantification (measurement: ex/em = 605/670 nm). Representative *ex vivo* fluorescence images and corresponding biodistribution quantification (n = 3, mean ± s.d.) of major organs resected from mice administered with (b, c) Ctrl (PBS) (measurement: ex/em = 465/570 nm).

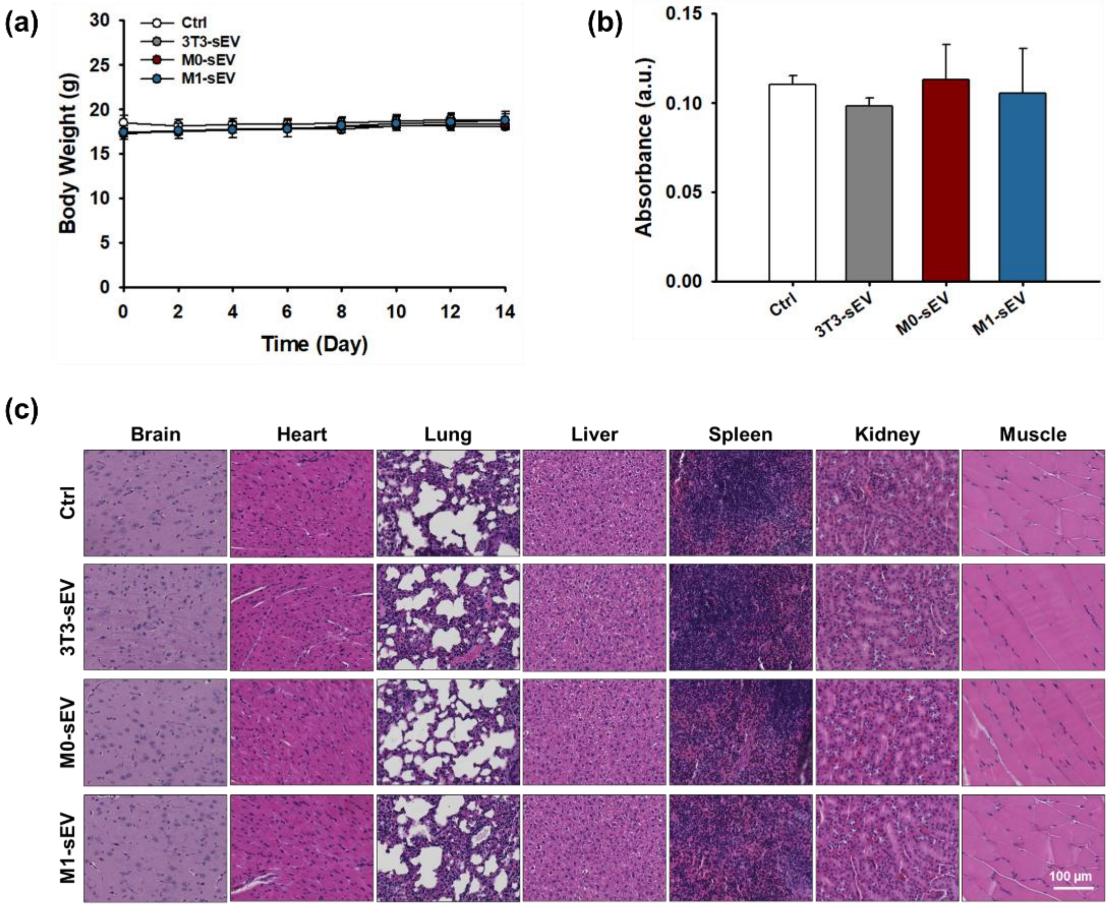


**Figure S4**. Safety evaluation of sEVs. (a) Body weight monitoring of mice following tail-vein i.v. injection. (b) Serum IL-6 levels measured by ELISA two weeks after the first injection (n = 3, mean ± s.d.). (c) H&E-stained histological images of major organs collected two weeks post-injection.

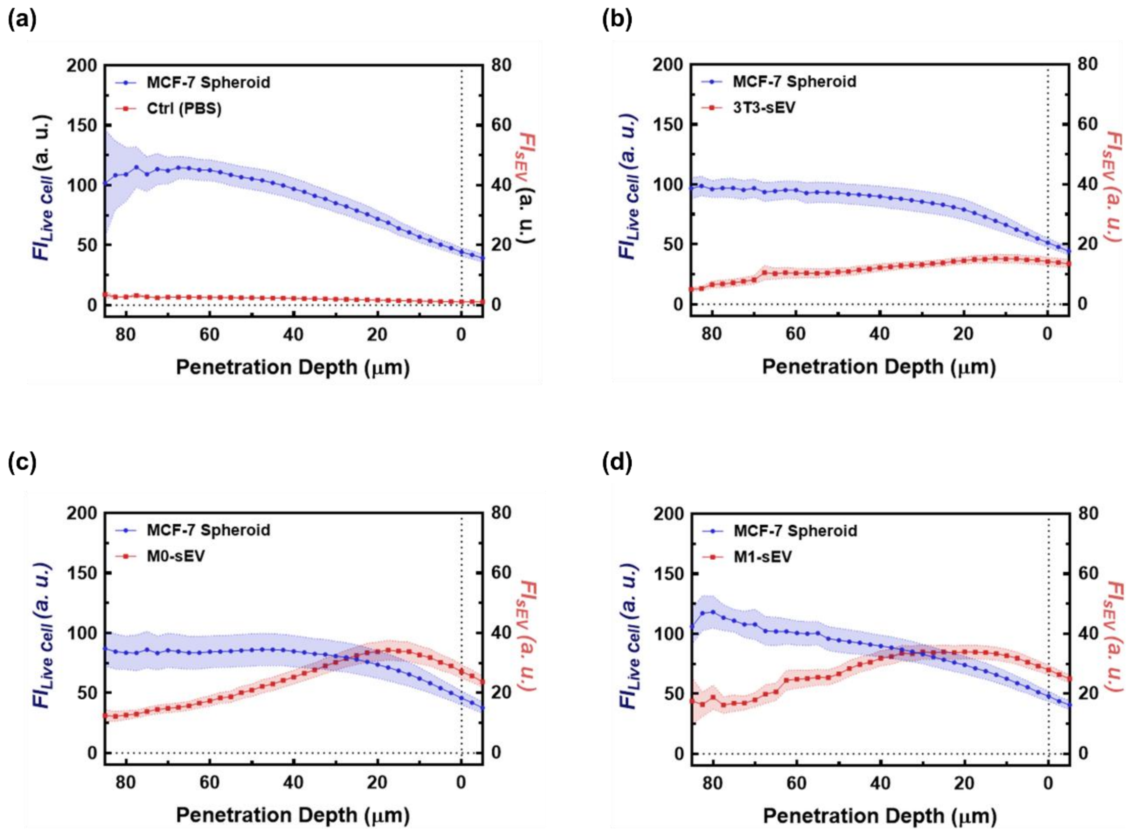


**Figure S5**. Quantification of fluorescence intensity in spheroid penetration depth for (a) Ctrl, (b) 3T3-sEV, (c) M0-sEV, and (d) M1-sEV (n > 30).

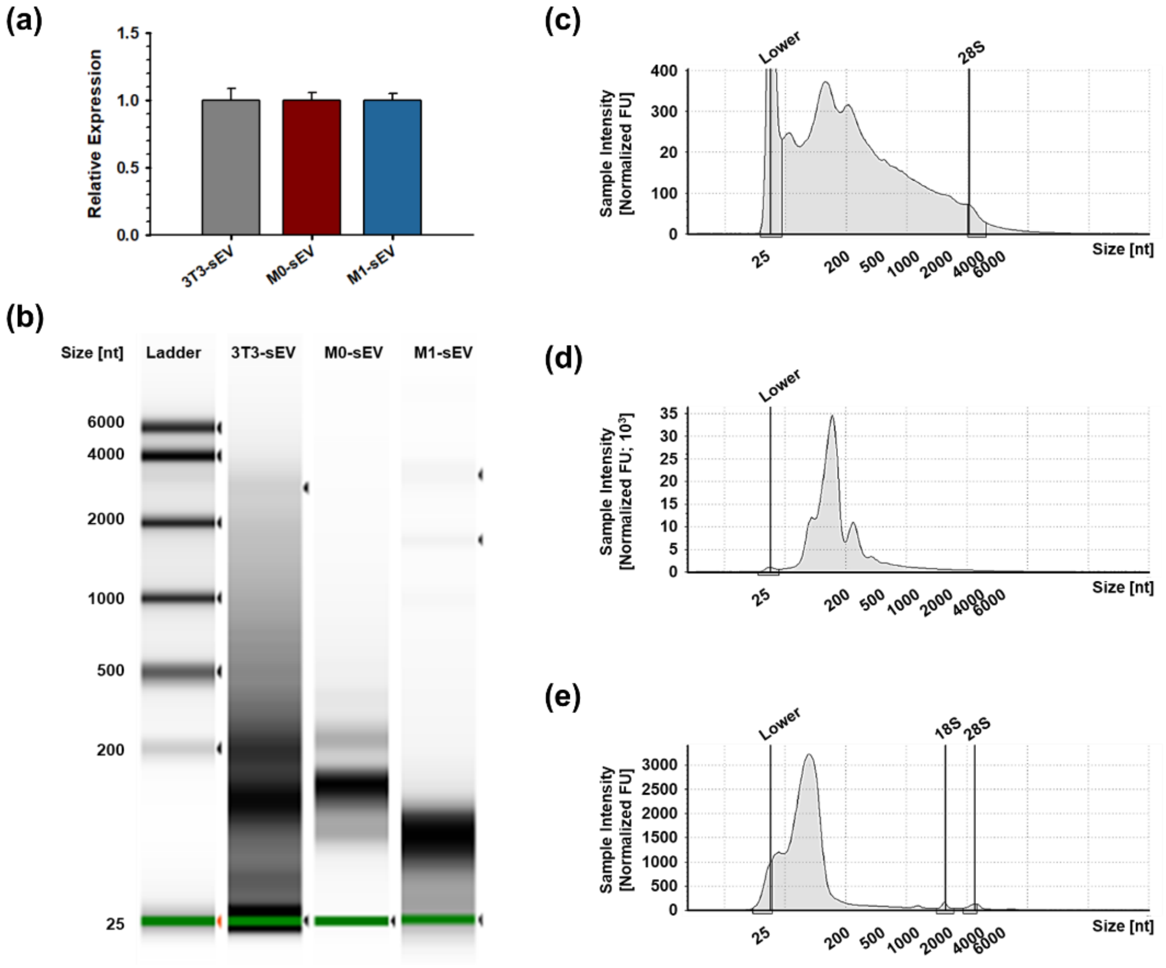


**Figure S6**. (a) Relative expression of U6 housekeeping gene measured by qRT-PCR. Quality control analysis of RNA extracted from sEVs: (b) Automated RNA electrophoresis using High Sensitivity RNA ScreenTape. Corresponding RNA profiles of (c) 3T3-sEV (Conc. 1,470 pg/mL), (d) M0-sEV (Conc. 32,300 pg/mL), and (e) M1-sEV (Conc. 5,050 pg/mL).

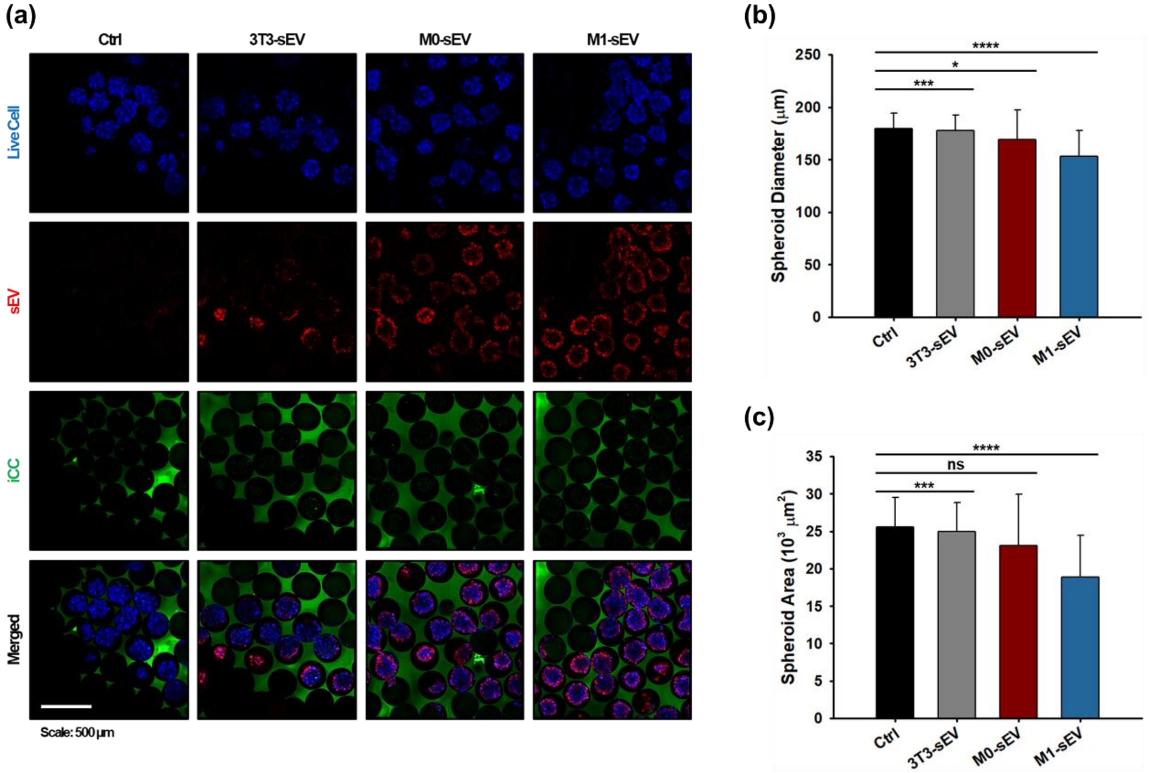


**Figure S7**. (a) TP-CLSM images of DiD-labeled sEV uptake in MCF-7 spheroids in iCC framework (Blue: live cells; Green: 5-DTAF-labeled iCC; Red: DiD-labeled sEV). Quantification of spheroid (b) diameter and (c) area from TP-CLSM images.

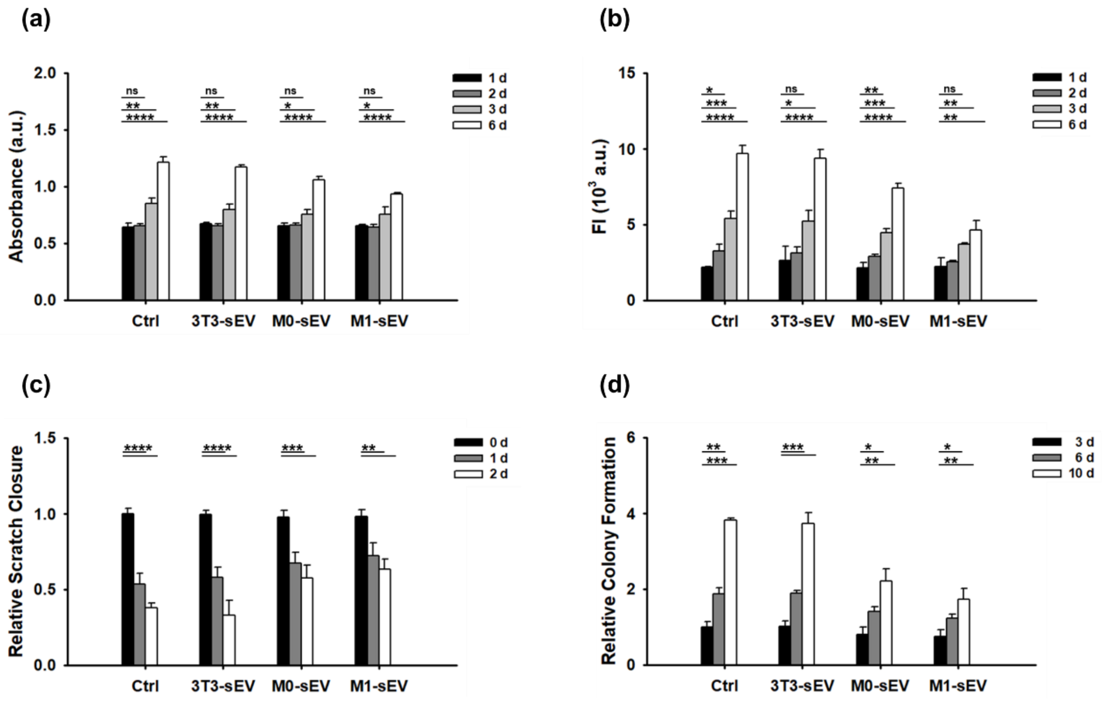


**Figure S8**. Quantification of time-dependent changes of anti-cancer effects within groups with statistical analysis. (a) WST-8 assay (n = 4, mean ± s.d.), (b) Resazurin assay (n = 4, mean ± s.d.), (c) Scratch assay (n = 4, mean ± s.d.), and (d) Clonogenic assay (n = 3, mean ± s.d.). One-way ANOVA with Tukey's post-test. ns: not significant; *p < 0.05; **p < 0.01; ***p < 0.001; ****p < 0.0001.

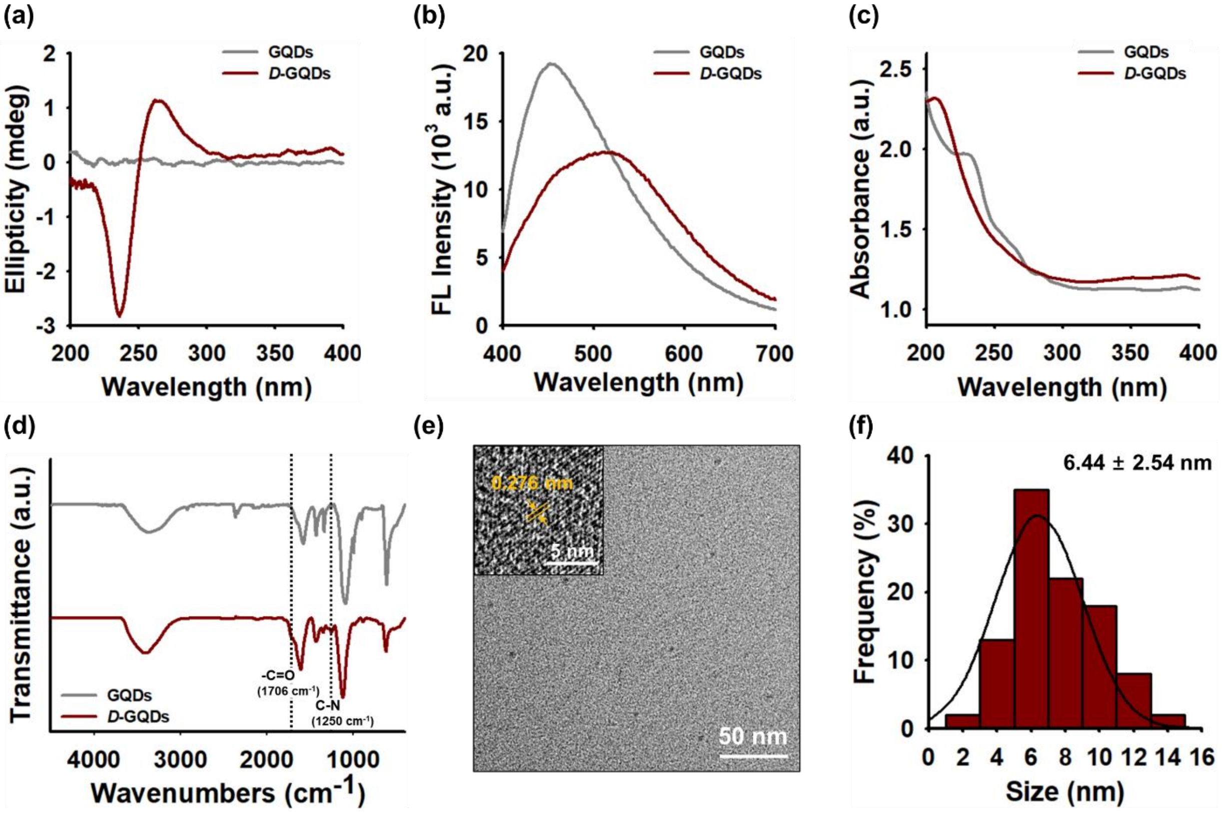


**Figure S9**. Characterization of chiral graphene quantum dots (GQDs). (a) Circular dichroism spectra. (b) Fluorescence spectra excited at 360 nm. (c) UV–vis absorption spectra. (d) FTIR spectra. (e) TEM image showing crystalline lattice structure of *D*-GQDs. (f) Size distribution of *D*-GQDs based on TEM image analysis (n = 100, mean ± s.d.).

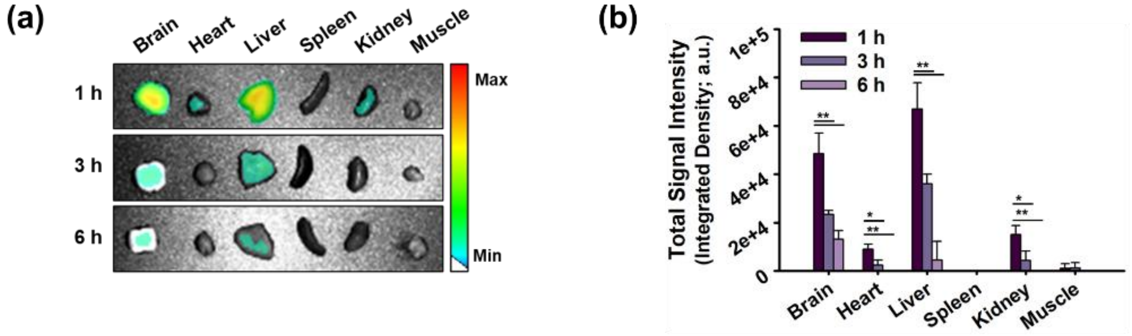


**Figure S10**. Representative *ex vivo* fluorescence images and corresponding biodistribution quantification (n = 3, mean ± s.d.) of major organs resected from mice administered with (a, b) *D*-GQDs (measurement: ex/em = 465/570 nm). One-way ANOVA with Tukey's post-test. ns: not significant; *$p < 0.05$; **$p < 0.01$; ***$p < 0.001$; ****$p < 0.0001$.

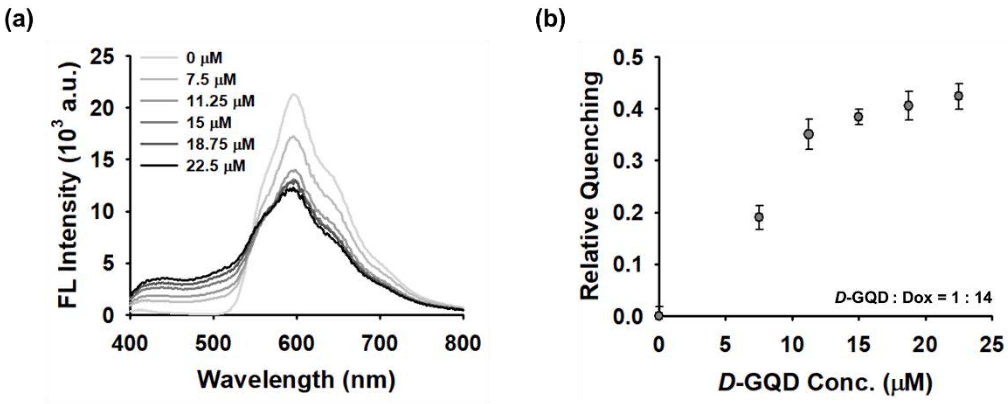


**Figure S11**. Characterization of Dox/*D*-GQD complex. (a) Fluorescence intensity and (b) relative quenching efficiency of Dox (200 μM) incubated with varying concentrations of *D*-GQDs (0–22.5 μM; n = 3, mean ± s.d.).

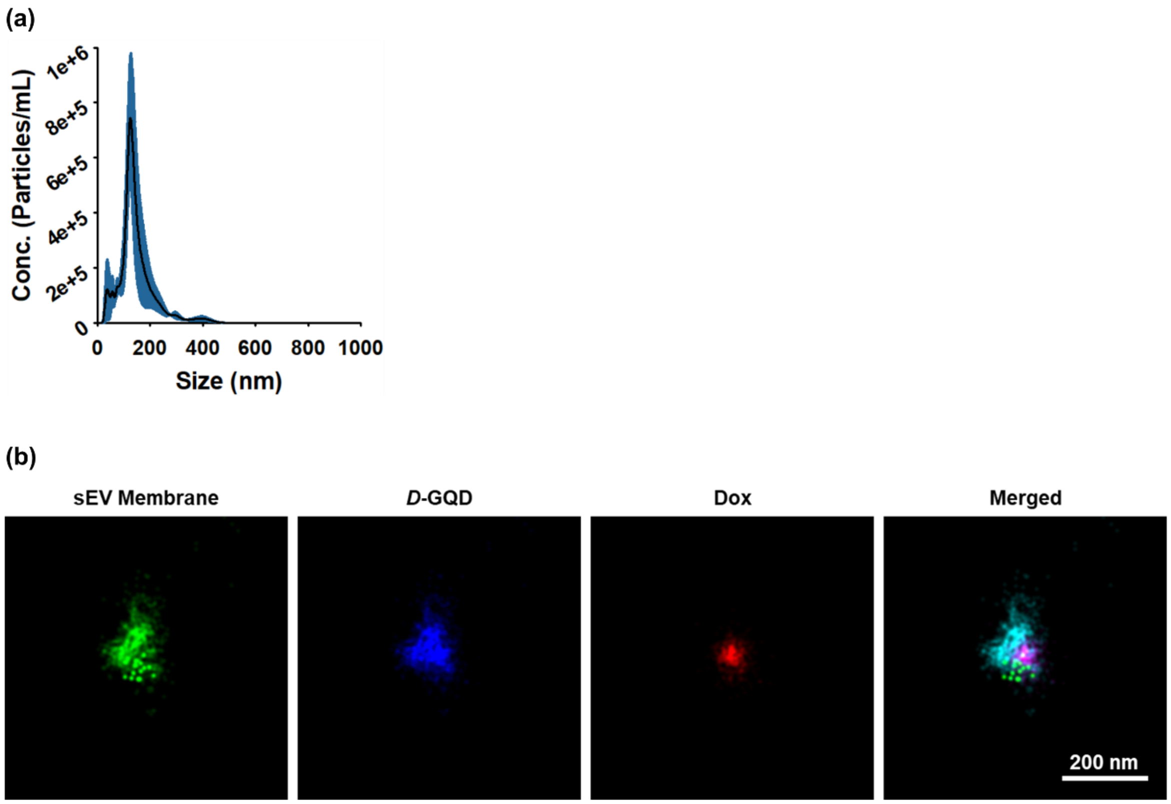


**Figure S12**. Characterization of siRNA-loaded sEVs via *D*-GQD. (a) NTA showing size distribution and particle concentration (n = 5). (b) Super-resolution images of individual sEVs obtained by single-molecule localization microscopy.

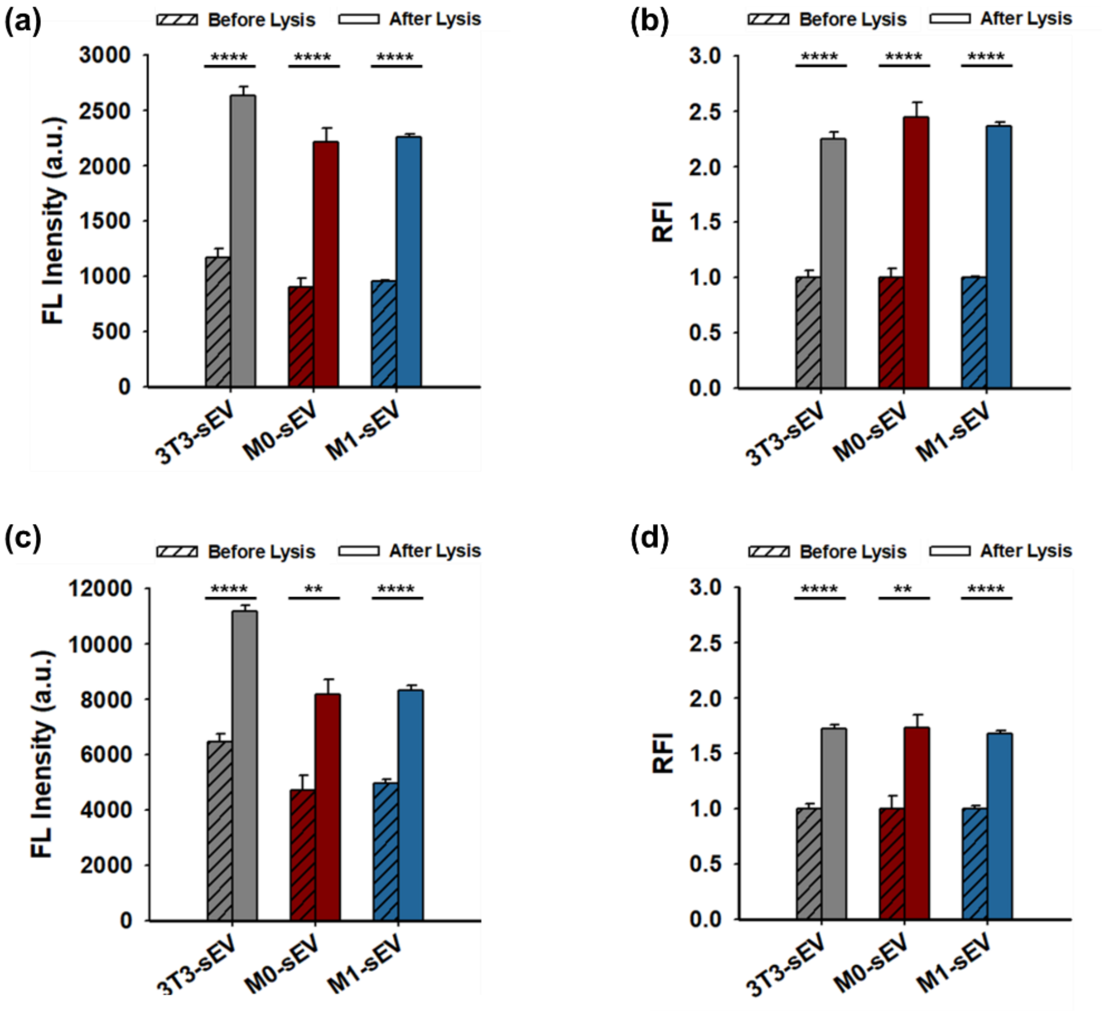


**Figure S13**. Fluorescence recovery test. Fluorescence intensity and relative fluorescence intensity (RFI) of (a, b) DiO (ex/em: 470/520 nm) on sEV membranes and (c, d) Dox (ex/em: 490/560 nm) encapsulated within sEVs, measured before and after membrane lysis with Tween-20 ($n = 3$, mean ± s.d.). One-way ANOVA with Tukey's post-test. ns: not significant; *$p < 0.05$; **$p < 0.01$; ***$p < 0.001$; ****$p < 0.0001$.

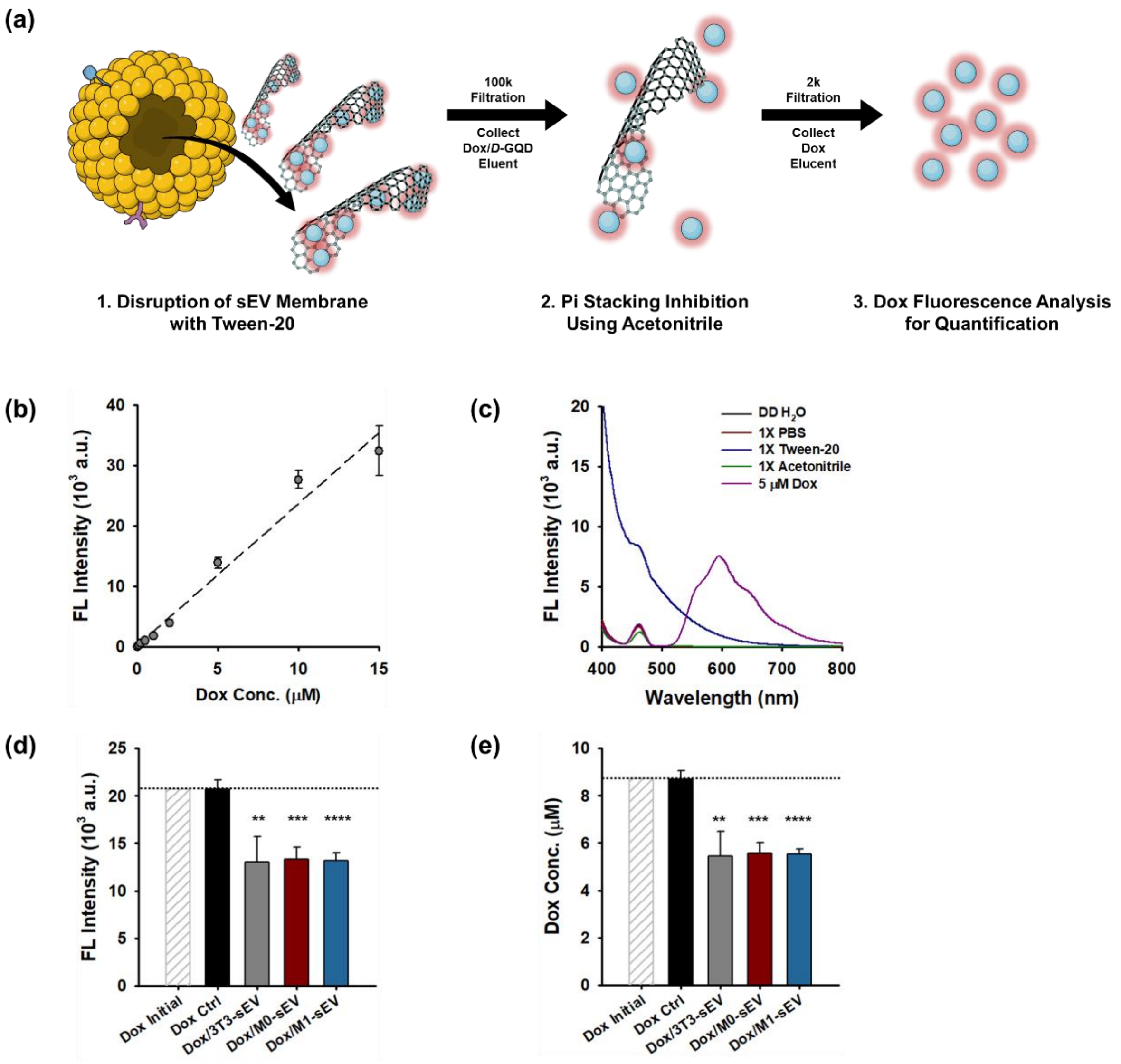


**Figure S14**. Quantification of Dox encapsulation efficiency in sEVs. (a) Schematic illustration of the quantification approach. (b) Calibration curve of Dox for determining the concentration released from sEVs. (c) Fluorescence spectra of solutions used throughout the process. (d) Fluorescence intensity of the final eluent containing Dox extracted from sEVs. (e) Quantification of Dox concentration based on the calibration curve and corresponding fluorescence intensity for each group ($n = 4$, mean ± s.d.). ex/em: 490/560 nm. One-way ANOVA with Tukey's post-test. ns: not significant; *$p < 0.05$; **$p < 0.01$; ***$p < 0.001$; ****$p < 0.0001$.

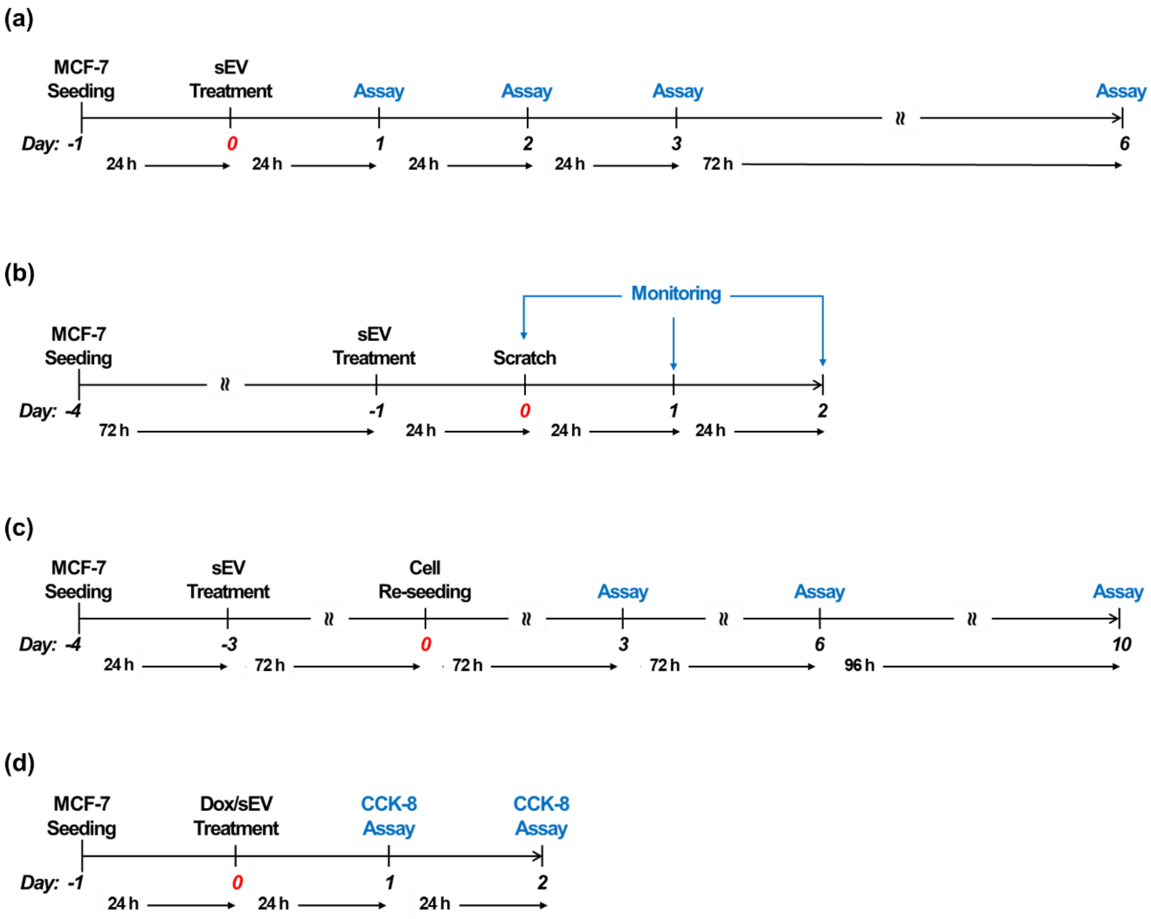


**Figure S15**. Workflow for *in vitro* assays. (a) WST-8 and resazurin assays for cell metabolic activity. (b) Scratch assay for cell migration. (c) Clonogenic assay for colony formation ability. (d) Cell viability assay for evaluating the therapeutic efficacy of Dox-loaded sEVs.

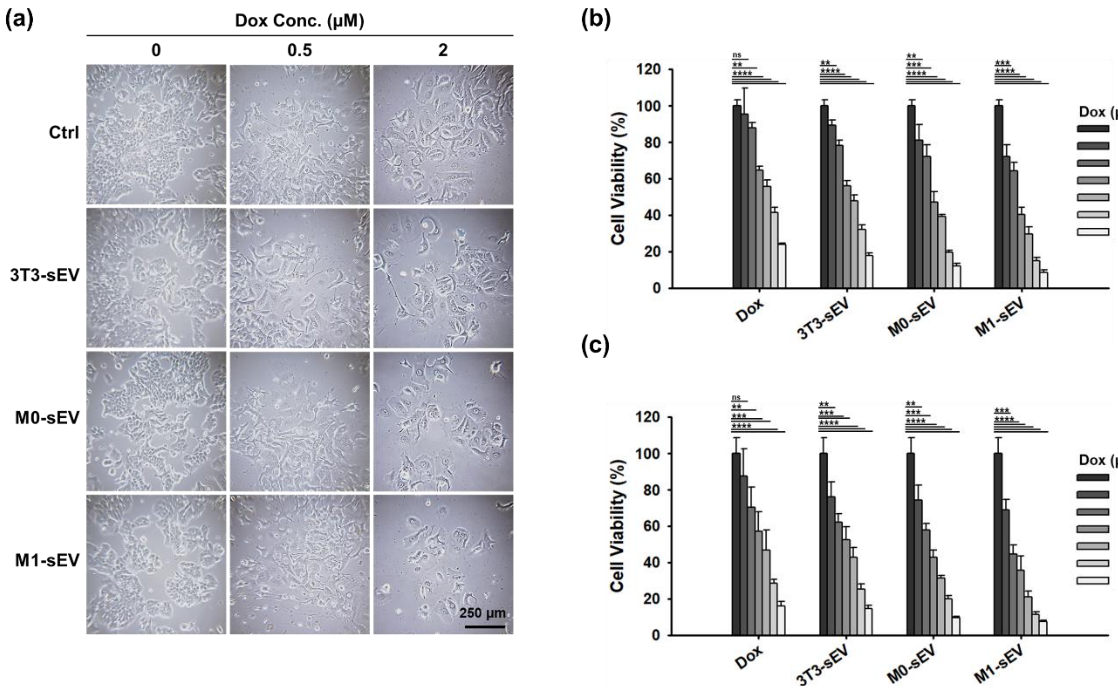


**Figure S16**. Quantification of time-dependent *in vitro* therapeutic effects within groups with statistical analysis. Cell viability of MCF-7 cells treated with Dox-loaded sEVs at various concentrations (0–10 μM) for (a, b) 24 h and (c) 48 h (n = 4, mean ± s.d.). One-way ANOVA with Tukey's post-test. ns: not significant; *p < 0.05; **p < 0.01; ***p < 0.001; ****p < 0.0001.

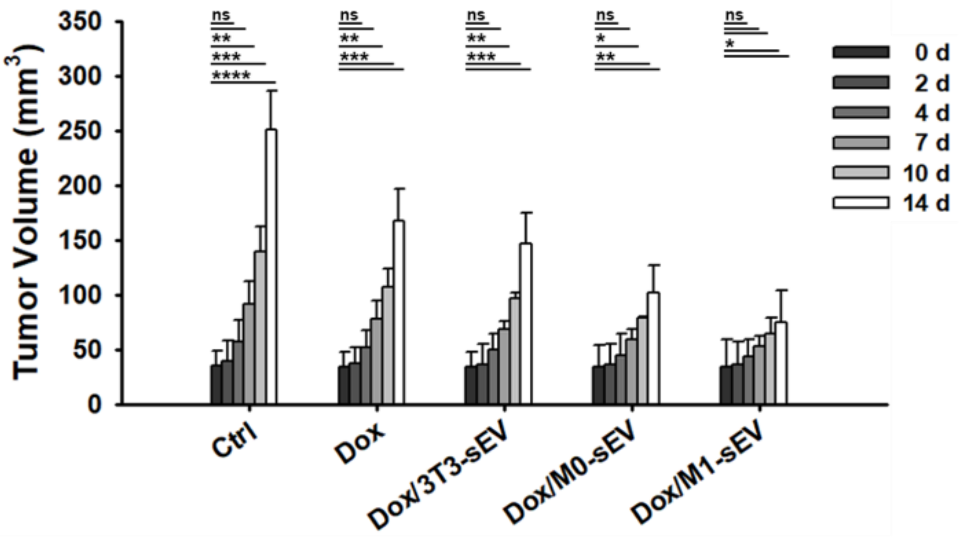


**Figure S17**. Quantification of time-dependent *in vivo* therapeutic effects within groups with statistical analysis. (n = 4, mean ± s.d.). One-way ANOVA with Tukey's post-test. ns: not significant; *$p < 0.05$; **$p < 0.01$; ***$p < 0.001$; ****$p < 0.0001$.

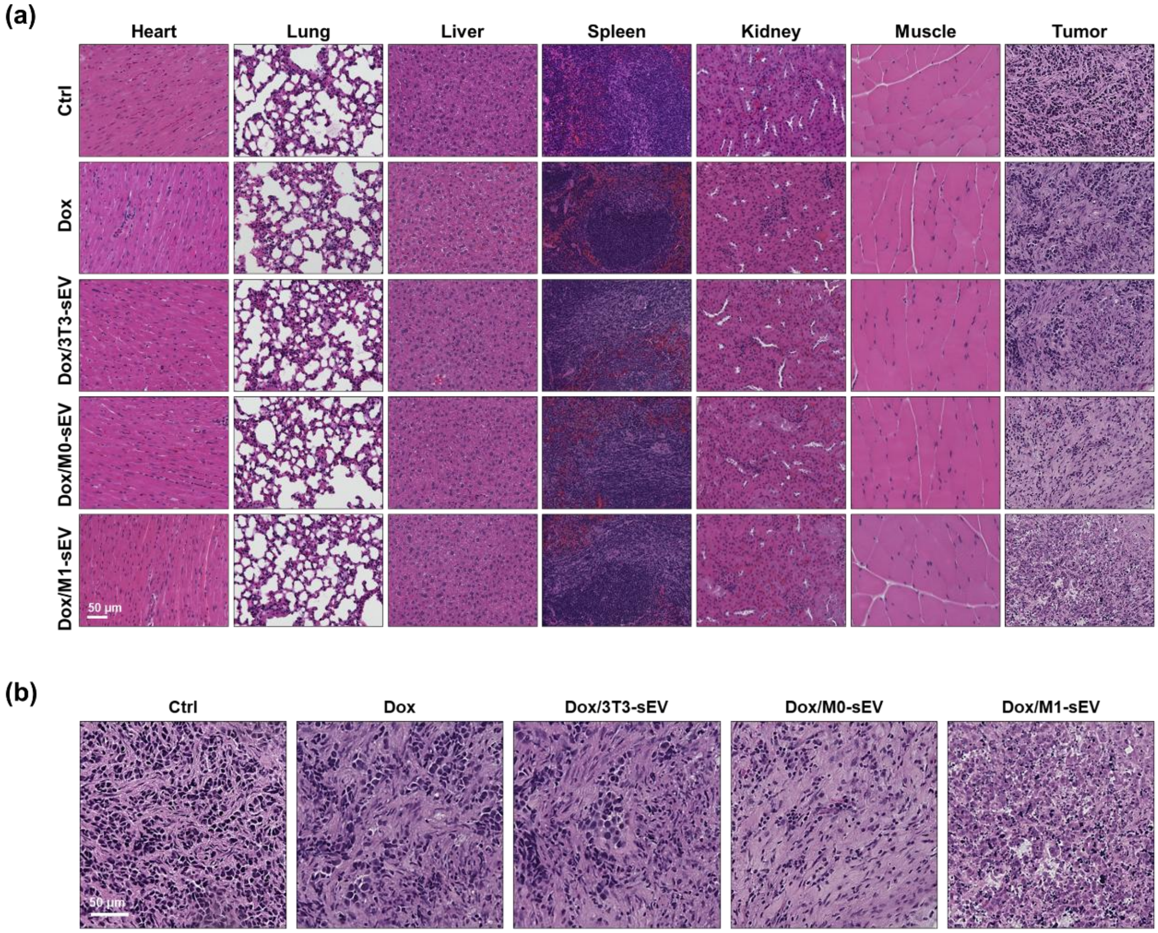


**Figure S18**. H&E-stained histological analysis. (a) Major organs and tumor tissues collected two weeks post-injection. (b) Magnified tumor images showing tissue damage.
